# The oxidative stress response of pathogenic *Leptospira* is controlled by two peroxide stress regulators which putatively cooperate in controlling virulence

**DOI:** 10.1101/2020.11.06.371039

**Authors:** Crispin Zavala-Alvarado, Samuel Garcia Huete, Antony T. Vincent, Odile Sismeiro, Rachel Legendre, Hugo Varet, Giovanni Bussotti, Céline Lorioux, Pierre Lechat, Jean-Yves Coppée, Frédéric J. Veyrier, Mathieu Picardeau, Nadia Benaroudj

**Author notes:** Corresponding author, (NB). Université de Paris, Sorbonne Paris Cité, Paris, France (Crispin Zavala-Alvarado, Samuel Garcia Huete). Current address: Institut Pasteur, Université de Paris, ^a^ Individualité Microbienne et Infection, ^b^ Biologie des Bactéries Pathogènes à Gram-Positif, ^c^ Biologie des ARN des Pathogènes Fongiques, F-75015 Paris, France.

## Abstract

Pathogenic *Leptospira* are the causative agents of leptospirosis, the most widespread zoonotic infectious disease. Leptospirosis is a potentially severe and life-threatening emerging disease with highest burden in sub-tropical areas and impoverished populations. Mechanisms allowing pathogenic *Leptospira* to survive inside a host and induce acute leptospirosis are not fully understood. The ability to resist deadly oxidants produced by the host during infection is pivotal for *Leptospira* virulence. We have previously shown that genes encoding defenses against oxidants in *L. interrogans* are repressed by PerRA (encoded by LIMLP_10155), a peroxide stress regulator of the Fur family. In this study, we describe the identification and characterization of another putative PerR-like regulator (LIMLP_05620) in *L. interrogans*. Protein sequence and phylogenetic analyses indicated that LIMLP_05620 displayed all the canonical PerR amino acid residues and is restricted to pathogenic *Leptospira* clades. We therefore named this PerR-like regulator PerRB. In *L. interrogans*, the PerRB regulon is distinct from that of PerRA. While a *perRA* mutant had a greater tolerance to peroxide, inactivating *perRB* led to a higher tolerance to superoxide, suggesting that these two regulators have a distinct function in the adaptation of *L. interrogans* to oxidative stress. The concomitant inactivation of *perRA* and *perRB* resulted in a higher tolerance to both peroxide and superoxide and, unlike the single mutants, a double *perRAperRB* mutant was avirulent. Interestingly, this correlated with major changes in gene and non-coding RNA expression. Notably, several virulence-associated genes (*clpB*, *ligA/B*, and *lvrAB*) were repressed. By obtaining a double mutant in a pathogenic *Leptospira* strain, our study has uncovered an interplay of two PerRs in the adaptation of *Leptospira* to oxidative stress with a putative role in virulence and pathogenicity, most likely through the transcriptional control of a complex regulatory network.

**Author summary:** Leptospirosis is a widespread infectious disease responsible for over one million of severe cases and 60 000 fatalities annually worldwide. This neglected and emerging disease has a worldwide distribution, but it mostly affects populations from developing countries in sub-tropical areas. The causative agents of leptospirosis are pathogenic bacterial *Leptospira* spp. There is a considerable deficit in our knowledge of these atypical bacteria, including their virulence mechanisms. In addition to the *Leptospira* PerRA regulator that represses defenses against peroxide, we have identified and characterized a second PerR regulator in pathogenic *Leptospira* species (PerRB) that participates in *Leptospira* tolerance to superoxide. Phenotypic and transcriptomic analyses of single PerRA and PerRB mutants suggest that the two PerRs fulfill distinct functions in the adaptation to oxidative stress. Concomitant inactivation of PerRA and PerRB resulted in a higher tolerance to both peroxide and superoxide. Moreover, the *perRAperRB* mutant lost its virulence. Major changes in gene expression, including a decreased expression of several virulence factors, were observed in the double *perRAperRB* mutant. Our study suggests that PerRA and PerRB cooperate to orchestrate a complex regulatory network involved in *Leptospira* virulence.

## Introduction

Pathogenic *Leptospira* spp. are aerobic diderm bacteria of the spirochetal phylum that are the causative agents of leptospirosis, the most widespread zoonosis [1]. More than one million cases of leptospirosis are currently estimated annually in the world, with about 59000 deaths [2]. This disease is considered as a health threat among impoverished populations in developing countries under tropical areas [2], but the number of reported cases of leptospirosis are also on the rise in developed countries under temperate climates [3]. Rodents are asymptomatic reservoir for leptospires as the bacteria colonize the proximal renal tubules of these animals. Leptospires are shed in the environment through the urine of infected animals and leptospirosis is transmitted to other animal species and humans mostly by exposure to contaminated soils and water. *Leptospira* penetrate an organism through abraded skins and mucous membranes, subsequently disseminate within the bloodstream and rapidly spread to multiple tissues and organs (including kidney, liver, lungs). Clinical manifestations range from a mild flu-like febrile state to more severe and fatal cases leading to hemorrhages and multiple organ failure. Because of the lack of efficient tools for genetic manipulation of *Leptospira* spp. and their fastidious growth in the laboratory conditions, our understanding of the mechanism of pathogenicity and virulence as well as the basic biology of these pathogens have been greatly hampered [4, 5].

During their life cycle, most bacteria will be exposed to reactive oxygen species (ROS), such as superoxide anion (O_2_) and hydrogen peroxide (H_2_O_2_), that are produced endogenously through the aerobic respiratory chain or encountered in the environment [6]. These ROS are also produced, together with hypochlorous acid (HOCl) and nitric oxide anion (NO), as powerful and efficient weapons by eukaryotic innate immune cells upon infection by pathogenic bacteria [7]. ROS cause oxidative damage to cellular components (proteins, DNA and lipids) and this would result in bacterial death if bacteria had not developed scavenging enzymes to counteract the deadly effect of ROS, including catalase, peroxidases and superoxide dismutase or reductase (SOD, SOR).

ROS production is increased in human and animals upon infection by *Leptospira* [8–10]. In fact, the ability to detoxify H2O2 is essential for *Leptospira* virulence as inactivation of the catalase-encoding gene led to virulence attenuation in *L. interrogans* [11]. The oxidative stress response in pathogenic *Leptospira* spp. is controlled by the ROS sensing transcription factor peroxide stress regulator (PerR) (encoded by LIMLP_10155/LIC12034/LA1857) [12, 13]. PerR belongs to the Fur (Ferric uptake regulator) transcriptional factor family. It is a metalloprotein that binds DNA in the presence of iron or manganese and represses the expression of catalase and peroxidases [14–16]. In the presence of peroxide, iron-bound PerR is oxidized on the histidine residues participating in iron coordination. Iron is released and PerR dissociates from DNA. As a consequence, peroxide-scavenging enzyme repression is alleviated [17, 18].

We have very recently characterized the transcriptional response to hydrogen peroxide in the pathogen *L. interrogans* and shown that these bacteria respond to sublethal H2O2 concentration by increasing the expression of catalase and of two peroxidases (Alkylperoxiredoxin (AhpC) and Cytochrome C peroxidase (CCP)) [19]. When *Leptospira* were exposed to deadly H2O2 concentration, additional enzymes with a putative role as antioxidants and/or in repair of oxidized cysteines in proteins were up-regulated, including several thiol oxidoreductases (thioredoxin, glutaredoxin, DsbD, and Bcp-like proteins) [19]. Several genes of the LexA regulon (*recA*, *recN*, *dinP*) and other genes with putative role in DNA repair (*mutS*, *radC*) had a higher expression as well as genes encoding for canonical chaperones (*dnaK/dnaJ/grpE*, *groES/EL*, and *hsp15/20*) [19]. Only genes coding for the catalase and peroxidases were under the control of PerR and our study has revealed a complex regulatory network independent of PerR involving other transcriptional regulators, sigma factors, two component systems and non-coding RNAs [19]. During the course of this study, we noticed that an ORF encoding a Fur-like regulator (LIMLP_05620/LIC11158/LA2887) was up-regulated when *Leptospira* were exposed to a deadly concentration of H2O2.

Here, we report the functional characterization of the LIMLP_05620-encoding Fur-like regulator (PerRB) in the adaptation of *Leptospira* to oxidative stress. *In silico* analyses demonstrate that PerRB exhibits PerR canonical amino acid residues and is the closest relative of PerRA (encoded by LIMLP_10155). Furthermore, a *perRB* mutant has a higher tolerance to superoxide. By obtaining a double mutant where *perRA* and *perRB* are concomitantly inactivated, we have also investigated the interplay between these two PerR-like regulators in the adaptation to oxidative stress and virulence of *L. interrogans*. Our study suggests that LIMLP_05620 encodes a second PerR-like regulator in pathogenic *Leptospira* species that putatively cooperates with PerRA in controlling *Leptospira* virulence.

## Results

### Identification of an ORF that encodes a novel putative PerR regulator in pathogenic *Leptospira* species

Regulators of the Fur family are homodimeric metalloproteins with a two-domain organization composed of an amino-terminal DNA binding domain and a carboxy-terminal dimerization domain (Fig 1A). The DNA binding domain contains a winged helix-turn-helix (HTH) DNA binding motif (H2-H4, in Fig1A) where the H4 helix mediates DNA binding. The dimerization domain consists of an *α*/*β* domain. The regulatory iron that controls DNA binding and dissociation is coordinated by histidine, aspartate, and glutamate residues located in a loop at the hinge of the amino and carboxy-terminal domains. Most of Fur-like regulators also coordinate a structural metal (zinc) through 2 cysteinate motifs (CxxC, where x designates any AA). This structural metal allows correct folding and dimeric assembly of the regulator.

**Fig 1.**
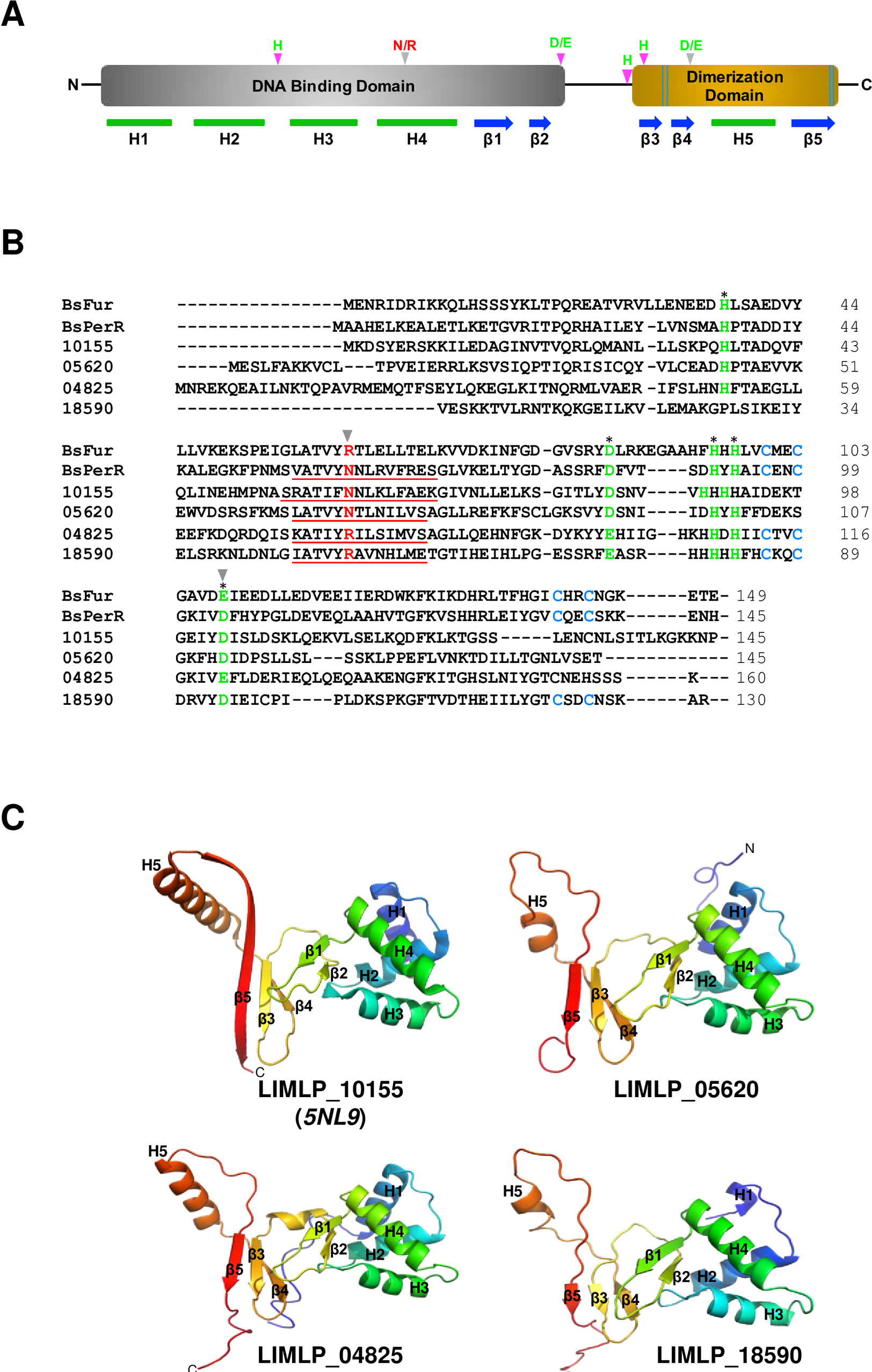
Analysis of the four Fur-like regulators of *L. interrogans*. (A) Schematic representation of the domain organization of a typical Fur-like regulator. The N-terminal DNA binding domain and the C-terminal dimerization domain are represented in grey and golden, respectively. The *α*-helix and *β*-strand secondary structures are indicated below in green and blue, respectively. The His, Asp and Glu residues involved in regulatory metal coordination are designated in green. The Arg/Asn residue involved in DNA binding specificity is marked in red. The Arg/Asn (involved in DNA binding specificity) and Asp/Glu residues (involved in H2O2 sensitivity) that allow distinguishing a Fur from a PerR are further emphasized with a grey arrow head. The two cysteinate motifs in the C-terminal domain involved in structural metal coordination are represented by the double blue lines in the C- terminal dimerization domain. (B) Comparison of the four Fur-like regulators of *L. interrogans* (LIMLP_10155, LIMLP_05620, LIMLP_04825, LIMLP_18590) with *B. subtilis* Fur and PerR. Primary sequence alignment was obtained by Clustal Omega (https://www.ebi.ac.uk/Tools/msa/clustalo/; [70]). The H4 DNA binding helix is underlined and the Arg/Asn residue involved in DNA binding specificity is designated in red. The residues of the regulatory metal coordination, including the Asp/Glu residue involved in H2O2 sensitivity, are marked in green and indicated with an asterisk. The Arg/Asn and Asp/Glu residues that allow distinguishing a Fur from a PerR are further emphasized with a grey arrow head. The cysteine residues of the structural metal coordination are marked in cyan. (C) Cartoon representation of the crystal structure of LIMLP_10155 (5NL9) and of the modeled structure of LIMLP_05620, LIMLP_04825 and LIMLP_18590. The modeled structures were obtained by searching homologies between LIMLP_05620, LIMLP_04825 and LIMLP_18590 and protein with known crystal structure using PHYRE2 (http://www.sbg.bio.ic.ac.uk/~phyre2/html/page.cgi?id=index; [71]). Secondary structures are numbered as in (A).

Due to high conservation in folding, metal coordination and similarity in the metal-induced conformational switch controlling DNA binding, it is difficult to distinguish members of the Fur family on the sole basis of their primary sequence. However, in *B. subtilis*, a single amino acid residue in the H4 helix of the amino-terminal DNA binding domain (N61 and R61 for *B. subtilis* PerR and Fur, respectively) allows PerR and Fur to discriminate between their respective DNA sequence targets (PerR and Fur boxes, respectively) [20] (Fig 1B). In addition, D104 residue in the carboxy-terminal domain is pivotal in the PerR sensitivity to H2O2. The corresponding residue in Fur is a glutamate and mutating this residue to aspartate leads to H2O2 sensitivity [21] (Fig 1B). Therefore, N61 and D104, which are well-conserved in other bacterial species, are considered as canonical PerR amino acid residues [20, 21].

*L. interrogans* genome encodes 4 ORFs that share homology with regulators of the Fur family. Sequence alignment of these ORFs with the Fur and PerR from *B. subtilis* shows a good conservation of residues involved in the regulatory metal coordination (Fig 1B). Interestingly, two of the 4 Fur-like ORFs of *L. interrogans*, LIMLP_10155 (LIC12034/LA1857 encoding a PerR) and LIMLP_05620 (LIC11158/LA2887), exhibit the canonical asparagine (N60 and N68, respectively) and aspartate (D103 and D112, respectively) residues of a typical PerR. The third ORF encoding a putative Fur-like regulator, LIMLP_04825 (LIC11006/LA3094), possesses the two Fur typical residues, R76 and E121, respectively. The fourth ORF encoding a putative Fur-like regulator, LIMLP_18590 (LIC20147/LB183), possesses the typical Fur arginine residue in its putative H4 DNA binding helix (R51) but a typical PerR aspartate residue in the carboxy-terminal domain (D94). Of note, LIMLP_18590 has a glutamate residue at the position 96. Fold prediction suggests that the three Fur regulators encoded by LIMLP_05620, LIMLP_04825 and LIMLP_18590 adopt the two-domain organization typical of the Fur family depicted in the crystal structure of LIMLP_10155 (Fig 1C).

The closest relative of the PerR-encoding LIMLP_10155 is LIMLP_05620 with about 26% of sequence identity, and LIMLP_04825 and LIMLP_18590 are closest relatives that share 20% of sequence identity. LIMLP_05620 shares about 27% identity with the well-characterized *B. subtilis* PerR. The putative H4 helix in LIMLP_05620 (Leu63-Ser75) is relatively well conserved with that of *B. subtilis* (Val56-Ser69) (Fig1B-C) and LIMLP_05620 displays a typical regulatory metal coordination site (His44-Asp93-His99-His101-Asp112). As the LIMLP_10155-encoded PerR, LIMLP_05620 lacks the cysteinate motif involved in structural metal coordination [13] (Fig 1B). In contrast, both LIMLP_04825 and LIMLP_18590 have one or two cysteinate motifs for structural metal coordination (C113xxC116 and C86xxC89- C122xxC125, respectively). Therefore, LIMLP_05620 encodes a putative PerR-like regulator closely related to the PerR-encoding LIMLP_10155 whereas LIMLP_04825 and LIMLP_18590 could encode other type of Fur-like regulators (Fur, Zur, or Nur). We therefore annotated the LIMLP_10155 and LIMLP_05620 ORFs as *perRA* and *perRB*, respectively.

### Phylogenetic analysis of PerRA and PerRB in *Leptospira* species

To get a better understanding of the evolutionary relationship of the four Fur-like regulators in pathogenic *Leptospira*, we undertook phylogenetic analyses by searching for homologous sequences of the PerRA (LIMLP_10155), PerRB (LIMLP_05620), LIMLP_18590 and LIMLP_04825 proteins among the representative genomes present in GenBank. This revealed a large phylogenetic distribution with several branches (Fig 2A). The sequences homologous to the LIMLP_04825 and LIMLP_18590 proteins form two distinct groups (red and orange, respectively) separated by a common ancestor. To get better definition of phylogenetic relationships of PerR-like homologues, we performed analysis with only a subset of sequence (Fig 2B). This phylogenetic analysis shows two separated groups composed of the sequences of PerRA (LIMLP_10155) and PerRB (LIMLP_05620) (see S1Fig for a more complete and detailed tree).

**Fig 2.**
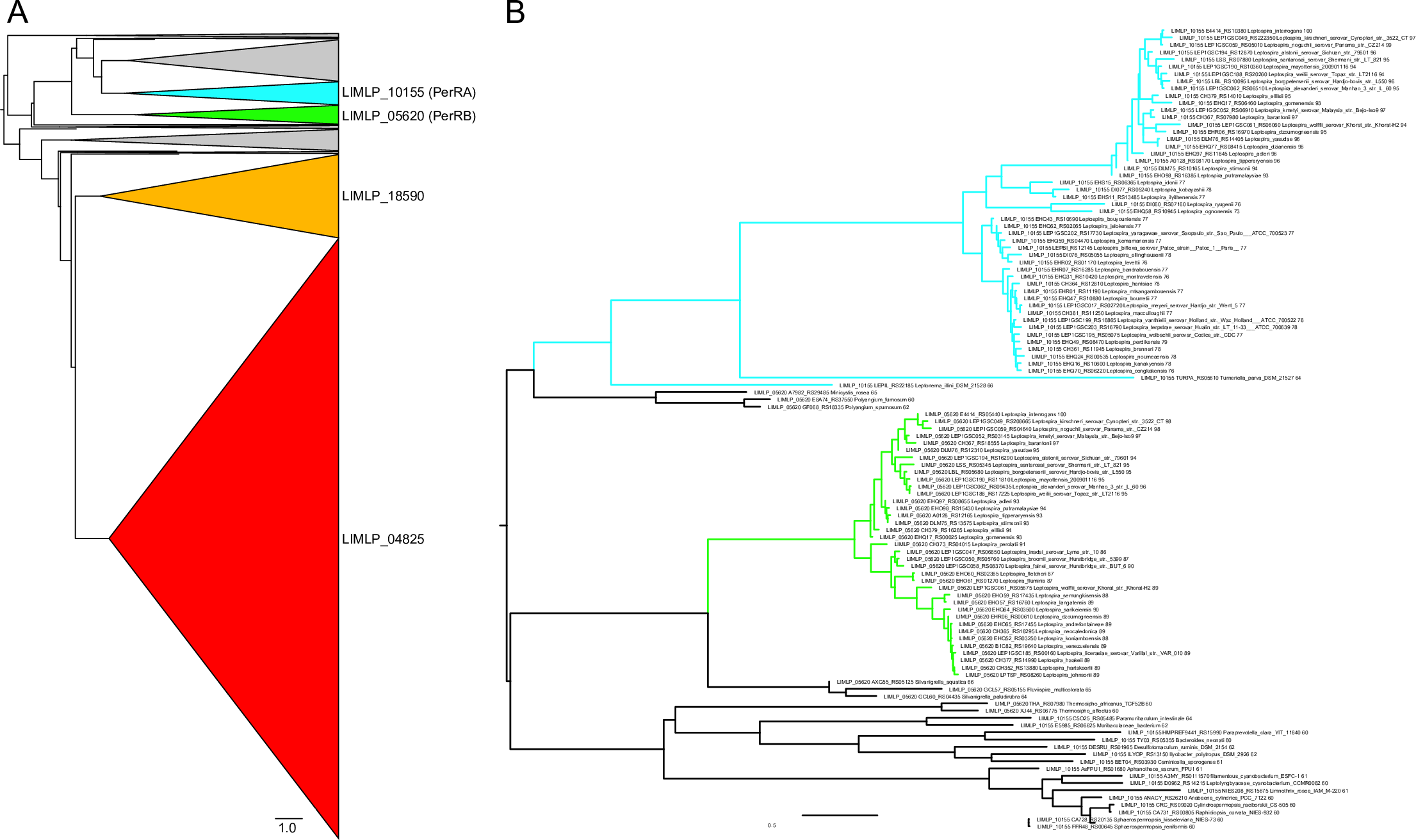
(A) Phylogenetic tree with a cartoon representation showing the distribution of the 1671 sequences putatively homologous to the PerRA (LIMLP_10155, cyan triangle), PerRB (LIMLP_05620, green triangle), LIMLP_18590 (yellow triangle) and LIMLP_04825 (red triangle) proteins. The gray triangles represent groups which are not monophyletic with the *Leptospira* sequences and which may therefore originate from other types of PerR or have had a species-specific evolution. (B) Phylogenetic tree showing the separation between PerRA (cyan) and PerRB (green).

The sequences of PerRA (LIMLP_10155) and PerRB (LIMLP_05620) ORFs from the strain *L. interrogans* serovar Manilae were searched and compared in all available genomes from the *Leptospira* genus (S1 Table). As seen in Fig 3, PerRA is present in the saprophytes S1 and S2 clades and in the P1 clade (highly virulent strains). This ORF is absent from most of the P2 clade (intermediate strains). However, there are two exceptions in the P2 clade species since homologues of PerRA are present in *L. dzoumogneensis* and *L. wolffii*. Additionally, this ORF is also present in other bacteria from the order *Leptospirales* such as *Turneriella parva* and *Leptonema illini* (Fig 2B). This suggests that PerRA was present in *Leptospirales* ancestor before *Leptospira* divergence and lost in the P2 clade. On the other side, PerRB ORF is only present in P1 and P2 clades and absent in all species from S1 and S2 clades (Fig 3). PerRB is also not found in other bacteria from the *Leptospirales* order (Fig 2B and S1 Fig). This restricted distribution suggests that the ancestor of pathogenic strains (P1 and P2 clades) has likely acquired PerRB after their divergence with other *Leptospira*. Overall, both PerRA and PerRB ORFs only coexist in P1 clade that comprises the highly virulent *Leptospira* species. Altogether, these findings indicate that pathogenic *Leptospira* strains encode a second putative PerR-like regulator that is absent in saprophytes.

**Fig 3.**
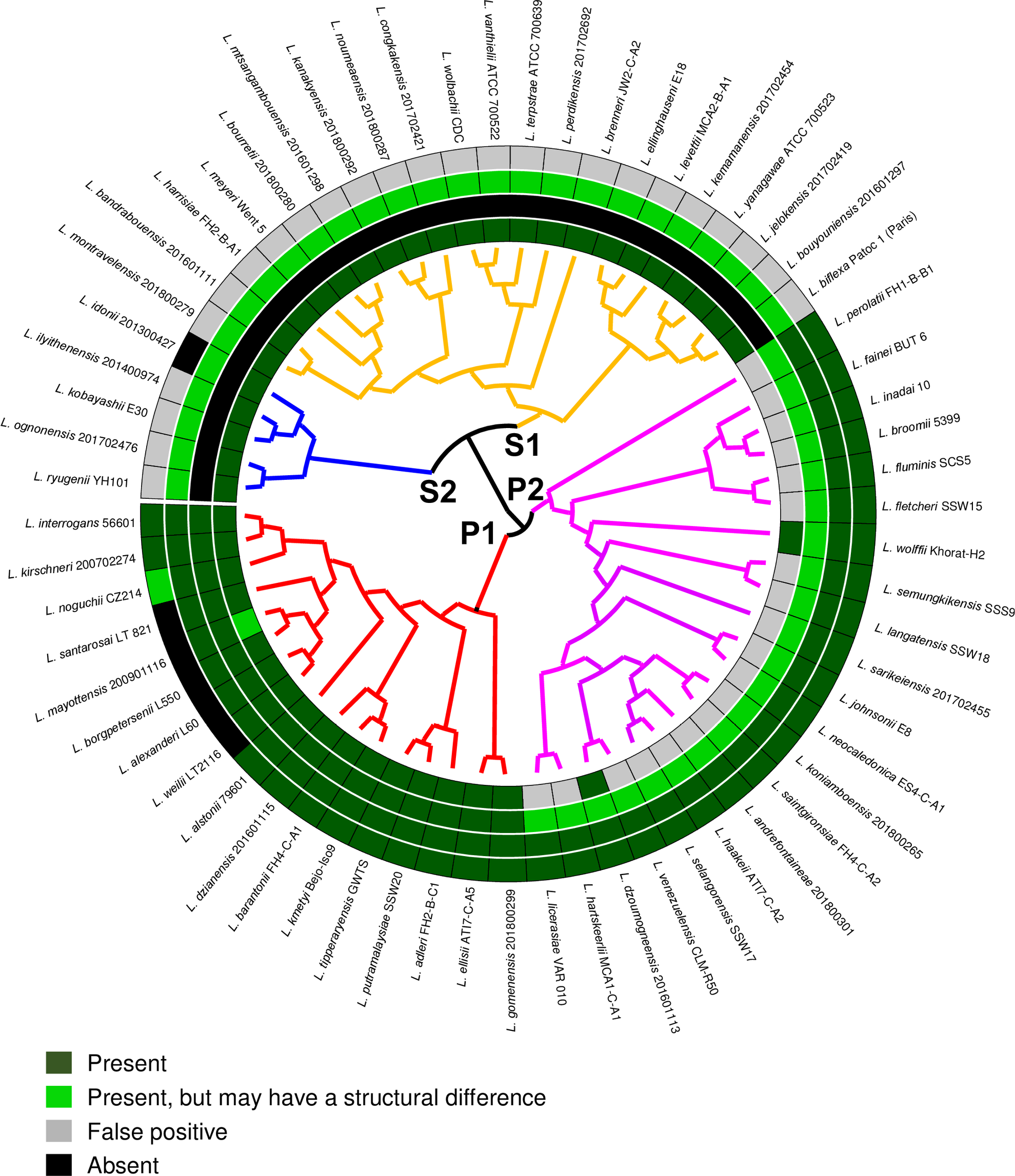
Distribution of the four Fur-like regulators of *L. interrogans* in the genus *Leptospira.* Circular phylogenetic tree with inner circles indicating the homology between each Fur-like regulator of *L. interrogans* with the closest homolog among representative genomes of *Leptospira* species. The branches are colored according to their classification into the four main subclades with P1 (highly pathogenic) in red, P2 (intermediates) in magenta, S1 (saprophytes) in yellow and S2 (new clade saprophytes) in blue [58]. The inner circles are, from the inside to the outside, PerRA (LIMLP_10155), PerRB (LIMLP_05620), LIMLP_04825 and LIMLP_18590. The green color gradient indicates the degree of homology (See S1 Table). Grey indicates the presence of a false positive (a different fur-like regulator with low homology) and black indicates the absence of orthologs.

### PerRB is involved in *L. interrogans* tolerance to ROS

As demonstrated previously [19], when *L. interrogans* are exposed to a subtlethal dose of H2O2 (10 µM for 30 min) *perRA* expression is increased by a 7-fold whereas that of *perRB* is unchanged (Fig 4). In the presence of a higher dose of H2O2 (1 mM for 1h), expression of both *perRA* and *perRB* was increased significantly by a 6-fold (Fig 4). Interestingly, the expression of the genes encoding the two other Fur-like regulators (LIMLP_04825 and LIMLP_18590) was not changed in the presence of H2O2. This indicates that PerRA and PerRB are the only Fur-like regulators that respond to H2O2 in *L. interrogans*.

**Fig 4.**
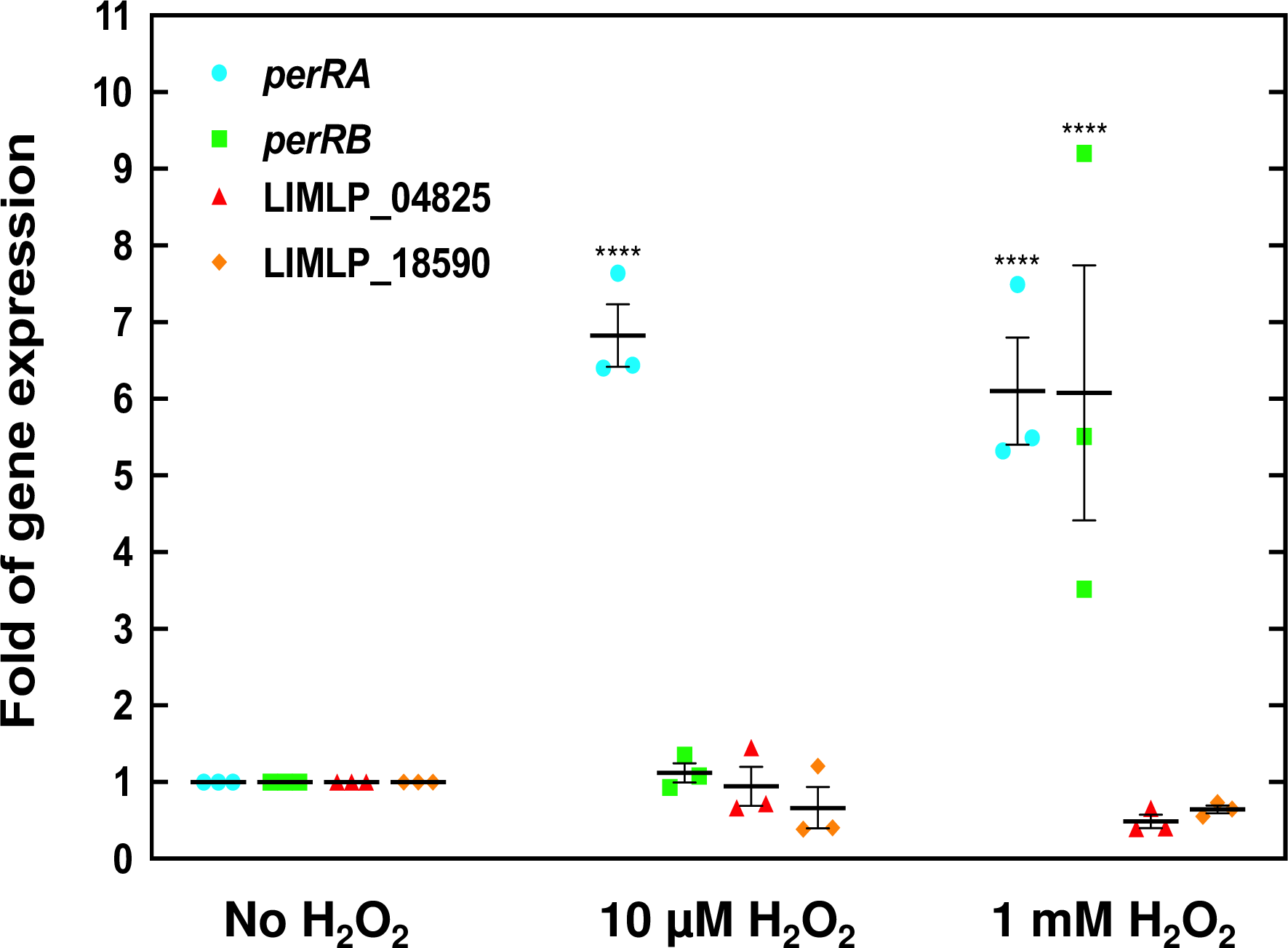
Increased *perRA* and *perRB* expression upon exposure to hydrogen peroxide. Exponentially growing *L. interrogans* were incubated in the absence or presence of 10 µM (for 30 min.) or 1 mM H2O2 (for 60 min.) and the expression of *perRA* (cyan circles), *perRB* (green squares), LIMLP_04825 (red triangles) and LIMLP_18590 (orange diamonds) was measured by RT-qPCR as described in the Material and Methods section using *flaB* as reference gene. Gene expression was normalized by that in untreated samples. Data are the means and standard errors of three independent biological replicates. Two-way Anova test indicated statistical significance in comparison with untreated samples (****, p- values<0.0001).

We have previously shown that inactivating *perRA* led to the derepression of *katE*, *ahpC* and *ccp* and to a higher tolerance to H2O2 [13, 19] (see S2 Fig). In addition, the *perRA* mutant exhibited a reduced ability to grow in the presence of the superoxide-generating compound paraquat [13]. A mutant with a transposon inserted into the PerRB-encoding LIMLP_05620 ORF was available in our random transposon mutant library and was used to investigate the role of PerRB in *L. interrogans* tolerance to ROS. Inactivating *perRB* did not have any effect on the ability of *L. interrogans* to tolerate deadly concentration of H2O2 (S2 Fig); however, it increases the capability of *Leptospira* to grow in the presence of paraquat (Fig 5B). Indeed, logarithmic growth of WT started after about 12 days in the presence of paraquat whereas that of the *perRB* mutant began after 5 days. Survival upon 1h exposure to 100 µM paraquat, as measured by colony-forming unit, showed that the *perRB* mutant had a 34-fold higher survival than that of the WT strain (Fig 5C). Resazurin reduction assay also confirmed the higher survival of the *perRB* mutant in the presence of paraquat (Fig 5D). Complementing in trans the *perRB* mutant (S3 Fig.) partially restored the WT growth and survival phenotypes in the presence of paraquat (Fig 5B-D). Indeed, logarithmic growth of the complemented strain started after 8-9 days and its survival decreased to a 9-fold higher value than that of the WT strain (Fig 5B-C). This indicates that PerRB is involved in *L. interrogans* tolerance to superoxide.

**Fig 5.**
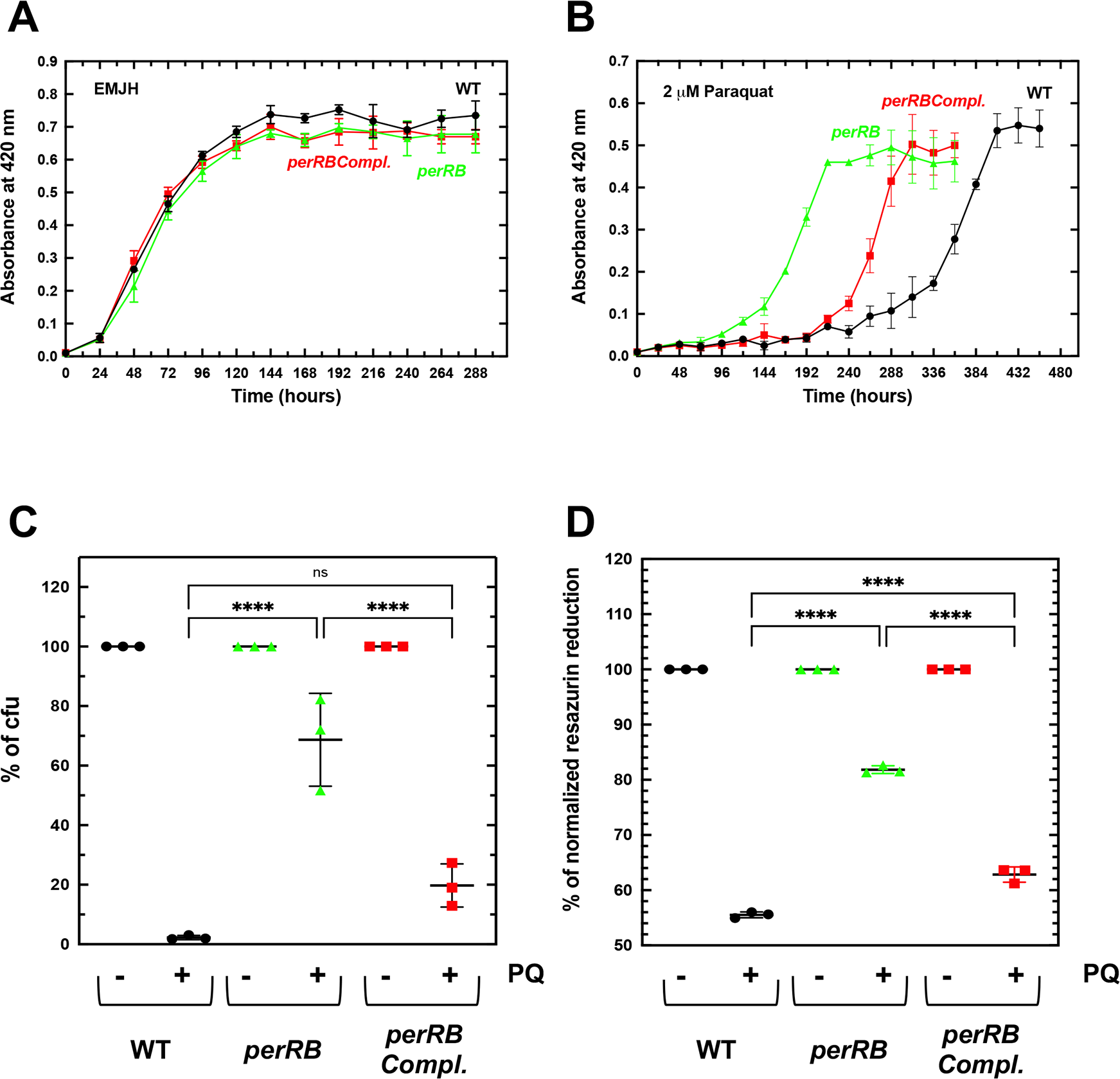
Effect of *perRB* inactivation on *Leptospira* growth and survival in the presence of superoxide-generating paraquat. *L. interrogans* WT containing the empty pMaORI vector (black circles), the *perRB* mutant containing the empty pMaORI vector (green triangles) or the *perRB* mutant containing the pMaORI vector expressing LIMLP_05620 (red squares) were cultivated in EMJH medium complemented with spectinomycin in the absence (A) or in the presence of 2 µM Paraquat (B). Growth was assessed by measure of absorbance at 420 nm. Data are means and standard errors of four independent biological replicates. In panel B, two-Anova test indicated that the differences between the *perRB* mutant and the WT were statistically significant from 144 to 360 h (p-value<0.0001), the differences between the *perRB* mutant and the complemented strain were statistically significant from 168 to 264 h (p-value<0.0001), and the differences between the complemented strain and the WT were statistically significant from 264 to 360 h (p-value<0.0001). Quantitative survival tests were performed by incubating exponentially growing WT containing the empty pMaORI vector (black circles), the *perRB* mutant containing the empty pMaORI vector (green triangles) or the *perRB* mutant containing the pMaORI vector expressing LIMLP_05620 (red squares) with 100 μM paraquat for 60 min. Percent of colony-forming unit (cfu) (C) and resazurin reduction (D) were determined as described in the Material and Method section. Results are shown as mean and SD of three independent experiments. Statistical significance was determined by a One-way Anova test (****, p- value<0.0001; ns, non significant).

### Identification of differentially-expressed genes upon *perRB* inactivation

To understand the role of PerRB in *L. interrogans* tolerance to ROS, we compared the global transcriptional profiles of the *perRB* mutant and WT strains. Differential gene expression analysis revealed changes in the transcription of 123 genes, with 59 and 64 down- and up-regulated, respectively (see S2 Table for a complete set of data). However, *perRB* inactivation did not lead to dramatic changes in gene expression as the majority of Log2FC (108 out of 123) ranged between -1 and 1 (S2 Table). These findings indicate that the absence of an active PerRB did not lead to substantial significant changes in genes expression when *Leptospira* are cultivated in the laboratory conditions (in EMJH medium at 30°C) and during the exponential phase.

Nevertheless, when examining the nature of the highest differentially-expressed genes in the *perRB* mutant, some tendencies could be observed. Many of the differentially-expressed ORFs were annotated as encoding proteins with unknown function and did not have homologs in the saprophyte *L. biflexa* strain (S2 Table and Table 1).

**Table 1.** Differentially-expressed ORFs upon perRB inactivation. Selected up-and down-regulated genes in the *perRB* mutant with an adjusted p-value cutoff of 0.05. ^a^ Gene numbering is according to Satou *et al.* [22]. Corresponding genes of *L. interrogans* serovar Lai strain 56601 and serovar Copenhageni Fiocruz strain L1-130 are indicated in parenthesis. ^b^ Closest ortholog in the saprophytes *L. biflexa* serovar Patoc strain Patoc1. The absence of synteny is indicated in italic. Genes that are down-regulated upon *perRA* inactivation as determined previously [19] are indicated in bold. Down (*) and up (**) -regulated genes upon exposure to lethal H2O2 dose as determined previously [19].

| ORF ID<sup>a</sup> | Gene | *L. biflexa*<sup>b</sup> | Function | Log<sub>2</sub>FC | Adjusted p-value |
| --- | --- | --- | --- | --- | --- |
| **Down-regulated genes** |  |  |  |  |  |
| LIMLP_02490* (LIC12988/LA0587) |  | *LEPBI_I0886* | Lipase putative extracellular lipoprotein | -1.608 | 6.11e-04 |
| LIMLP_02845 (LIC12920) |  |  | Hypothetical | -1.084 | 3.46e-02 |
| LIMLP_03640** (LIC12763/LA0865) |  |  | Hypothetical | -1.102 | 1.06e-02 |
| LIMLP_03790* (LIC12736/LA0905) |  |  | Hypothetical | -1.244 | 1.42e-05 |
| **LIMLP_04255 (LIC10892/LA3244)** | *exbB* | LEPBI_I0149 | Biopolymer transport protein ExbB/TolQ | -0.947 | 1.07e-02 |
| LIMLP_06190** (LIC11265/LA2751) |  | LEPBI_I3113 | Disulfide oxidoreductase | -0.723 | 6.96e-03 |
| LIMLP_11400** (LIC12297/LA1456) | *radC* |  | DNA repair protein RadC | -0.619 | 1.17e-02 |
| LIMLP_13165** (LIC12631/LA1029) | *sph2* |  | Sphingomyelinase C | -1.152 | 8.10e-03 |
| LIMLP_14595* (LIC10628/LA3571) |  | LEPBI_I2694 | Cytochrome oxidase CcoP subunit | -0.583 | 3.40e-02 |
| LIMLP_14715** (LIC10606/LA3598) | *dps* | *LEPBI_I2540* | DNA-binding stress protein Dps | -0.896 | 1.61e-03 |
| **LIMLP_15470\*\* (LIC10454/LA3793)** |  | LEPBI_I0671 | Hemolysin (N-acyltransferase domain) | -1.317 | 3.40e-03 |
| LIMLP_15890 (LIC10377/LA0430) |  |  | Hypothetical putative lipoprotein | -1.353 | 3.90e-10 |
| LIMLP_16695 (LEPIC3326/LA4096) |  |  | Hypothetical | -1.124 | 8.14e-04 |
| **LIMLP_18235\*\* (LIC20078/LB099)** |  |  | Hypothetical | -1.098 | 1.69e-02 |
| **Up-regulated genes** |  |  |  |  |  |
| LIMLP_02880* (LIC12912/LA0688) | *cas5* |  | CRISPR-associated protein Cas5 | 0.881 | 5.84e-03 |
| LIMLP_02885* (LIC12911-12910/LA0689-0690) | *cas3* |  | CRISPR-associated protein Cas3 | 0.834 | 4.51e-03 |
| LIMLP_04075** (LIC12680/LA0974) |  |  | Adhesin/FimH-like protein/DuF1566 domain | 1.110 | 4.29e-03 |
| LIMLP_05450* (LIC11125/LA2933) |  |  | Diguanylate cyclase | 0.983 | 2.12e-04 |
| LIMLP_05455* (LIC11126/LA2932) |  |  | Diguanylate cyclase | 0.757 | 1.81e-02 |
| LIMLP_05460* (LIC11127/LA2930) |  |  | Diguanylate cyclase | 0.922 | 1.11e-03 |
| LIMLP_05480 (LA2928) |  |  | Hypothetical | 0.968 | 2.67e-02 |
| LIMLP_05485** (LIC11131/LA2926) |  |  | Diguanylate cyclase | 0.679 | 8.14e-04 |
| LIMLP_05845 (LIC11203/LA2827) |  |  | Diguanylate phosphodiesterase | 0.608 | 3.85e-02 |
| LIMLP_09580 (LIC11921/LA1980) |  | LEPBI_I1269 | Diguanylate phosphodiesterase | 0.547 | 1.61e-02 |
| LIMLP_17875 (LIC20015/LB017) | *hemN* | *LEPBI_I1166* | Coproporphyrinogen III oxidase HemN | 0.727 | 9.53e-03 |
| LIMLP_18375** (LIC20106/LB133) |  |  | Diguanylate phosphodiesterase | 0.593 | 1.99e-02 |
| LIMLP_18755 (LIC20176/LB225) |  |  | Hypothetical putative lipoprotein | 1.453 | 3.77e-04 |
| LIMLP_18760 (LIC20177/LB226) |  |  | Adhesin/FimH-like protein/ DUF1566 domain-containing protein | 1.095 | 3.39e-02 |
| LIMLP_19320* (LA1770) |  |  | AraC family transcriptional regulator | 1.279 | 4.51e-03 |
| LIMLP_19325* (LA1771) |  |  | Hypothetical | 1.262 | 4.05e-02 |

Several genes involved in the metabolism of c-di GMP were differentially-expressed upon *perRB* inactivation. C-di GMP is a secondary messenger in bacteria that regulates a variety of processes such as biofilm formation, motility, stress adaptation, and virulence. C-di GMP synthesis is catalyzed by diguanylate cyclases (DGCs) whereas its hydrolysis is catalyzed by phosphodiesterases (PDEs). DGCs and PDEs are numerous in pathogenic *Leptospira*, suggesting that c-di GMP fulfills an important role in sensing environmental signals when *Leptospira* infect and colonize a host. C-di GMP has been recently shown to regulate biofilm formation, motility and protection against environmental stress in pathogenic *Leptospira* [23]. Four DGCs (LIMLP_05450, LIMLP_05455, LIMLP_05460, LIMLP_05485) were up-regulated upon *perRB* inactivation (Table 1). These DGC-encoding ORFs are located in a gene cluster (LIMLP_05485-05450) that contains 7 ORFs coding for DGCs. LIMLP_05450, LIMLP_05455, LIMLP_05460, and LIMLP_05485 display the typical diguanylate cyclase GGDEF and sensory PAS domains. A DGC activity was demonstrated *in vitro* for LIMLP_05450, LIMLP_05455, LIMLP_05460 [24]. Three PDE-encoding ORFs (LIMLP_05845, LIMLP_9580, and LIMLP_18375) were also up-regulated in the *perRB* mutant.

Among the highest up-regulated genes, two ORFs (LIMLP_04075 and LIMLP_18760) encoded lipoproteins with a putative adhesin function. These proteins contain DUF1566 domain repeats which is also share by Lsa25, a Leptospiral surface adhesin that binds extracellular matrix (ECM) [25].

An ORF encoding an AraC transcriptional regulator (LIMLP_19320), and two ORFs of unknown function (LIMLP_18755 and 19325) were also among the most up-regulated. The orthologs of LIMLP_19320 and LIMLP_19325 in *L. interrogans* serovar Lai belongs to a genomic island (Lai GI B, LA1747-1851) that can excise from the chromosome and form an independent replicative plasmid [26, 27].

LIMLP_13165 was among the most down-regulated ORFs when *perRB* was inactivated. It encodes a secreted protein with sphingomyelinase C and hemolytic activities [28]. Another significantly down-regulated ORF encoded a protein with an acyl CoA acetyl tranferase domain annotated as a putative hemolysin (LIMLP_15470). This ORF is up-regulated when *L. interrogans* is cultivated in DMC implanted in rats [29] .

Among the down-regulated genes, several ORFs encode factors related to oxidative stress. LIMLP_04255 is part of a gene cluster (LIMLP_04240-04285) which code for a putative TonB-dependent transport system repressed by PerRA. We have previously shown that some genes of this cluster (LIMLP_04245, LIMLP_04270 and LIMLP_04280) are involved in *L. interrogans* tolerance to superoxide [19]. LIMLP_11400 encodes the DNA repair protein RadC and LIMLP_14715 is a homolog of the *E. coli* Dps, a protein that sequesters iron and protects DNA from oxidative damage. LIMLP_06190 encodes a putative disulfide oxidoreductase with the N-terminal ScdA domain (DUF1858). In *S. aureus*, ScdA is a di-iron protein involved in repair of oxidatively damaged iron-sulfur cluster proteins [30]. LIMLP_14595 encodes a putative transmembrane lipoprotein with a cytochrome-like domain that shows homology with the CcoP subunit of the cytochrome C oxidase and could function in the respiratory chain or be an enzyme cofactor.

Only 7 out of the 123 differentially-expressed genes in the *perRB* mutant were also differentially-expressed upon *perRA* inactivation with a similar inclination (S4 Fig) [19]. LIMLP_02010 and LIMLP_04325 were up-regulated whereas LIMLP_04255, LIMLP_11810, LIMLP_14225, LIMLP_15470 and LIMLP_18235 were down-regulated in the two mutants.

Notably, 82 out of the 123 differentially-expressed ORFs in the *perRB* mutant were also differentially-expressed upon exposure of *L. interrogans* to H2O2 (S4 Fig) [19]. Thus, 66% of the PerRB regulon is also regulated by the presence of H2O2. Interestingly, the majority of ORFs down-regulated in the *perRB* mutant, including the RadC and the Dps-encoding ORFs, were up-regulated in the presence of H2O2 (with Log2FCs of 3.46 and 1.10, respectively) (S4 Fig and Table 1). In contrast, many up-regulated ORFs in the *perRB* mutant had a lower expression in the presence of H2O2. For instance, the ORFs that code for Cas5 (LIMLP_02880), Cas3 (LIMLP_02885), and two DGCs (LIMLP_05450 and LIMLP_05455) were down-regulated upon exposure to H2O2 with Log2FCs lower than -1.21 (S4 Fig and Table 1).

### Concomitant inactivation of *perRA* and *perRB* leads to a higher resistance to ROS and to a lower virulence

In order to investigate whether PerRA and PerRB cooperate in regulating the adaptive response to ROS, we inactivated *perRA* by allelic exchange in the *perRB* mutant (S5 Fig). This allowed obtaining a double *perRAperRB* mutant in *L. interrogans*.

The double *perRAperRB* mutant had a growth rate comparable to that of the single *perRA* and *perRB* mutants and WT strains when *L. interrogans* were cultivated in EMJH medium (Fig 6A). However, entry in exponential phase was delayed if the culture medium was inoculated with stationary phase-adapted *perRAperRB* mutant (S5 Fig). We had already shown that a *perRA* mutant had a higher ability to grow and survive in the presence of deadly concentration of H2O2 but a slower growth in the presence of the superoxide-generating paraquat ([13], and see in S2 Fig and Figs 6B-C). In contrast, inactivating *perRB* led to a higher resistance to paraquat (Fig 5 and Fig 6C). In the presence of H2O2, the double *perRAperRB* mutant was able to grow with a rate comparable to that of the *perRA* mutant. In this condition, the *perRB* mutant and WT strains did not exhibit any growth (Fig6B). In the presence of paraquat, the double *perRAperRB* and the *perRB* mutants entered in logarithmic growth earlier than the WT strain (Fig 6C). Survival tests showed that the *perRAperRB* mutant had a higher survival than the WT upon exposure to H2O2 (Fig 6D) or to paraquat (Fig 6E). Therefore, the double *perRAperRB* mutant exhibited cumulative phenotypes of the respective single *perRA* and *perRB* mutants when *L. interrogans* are exposed to ROS.

**Fig 6.**
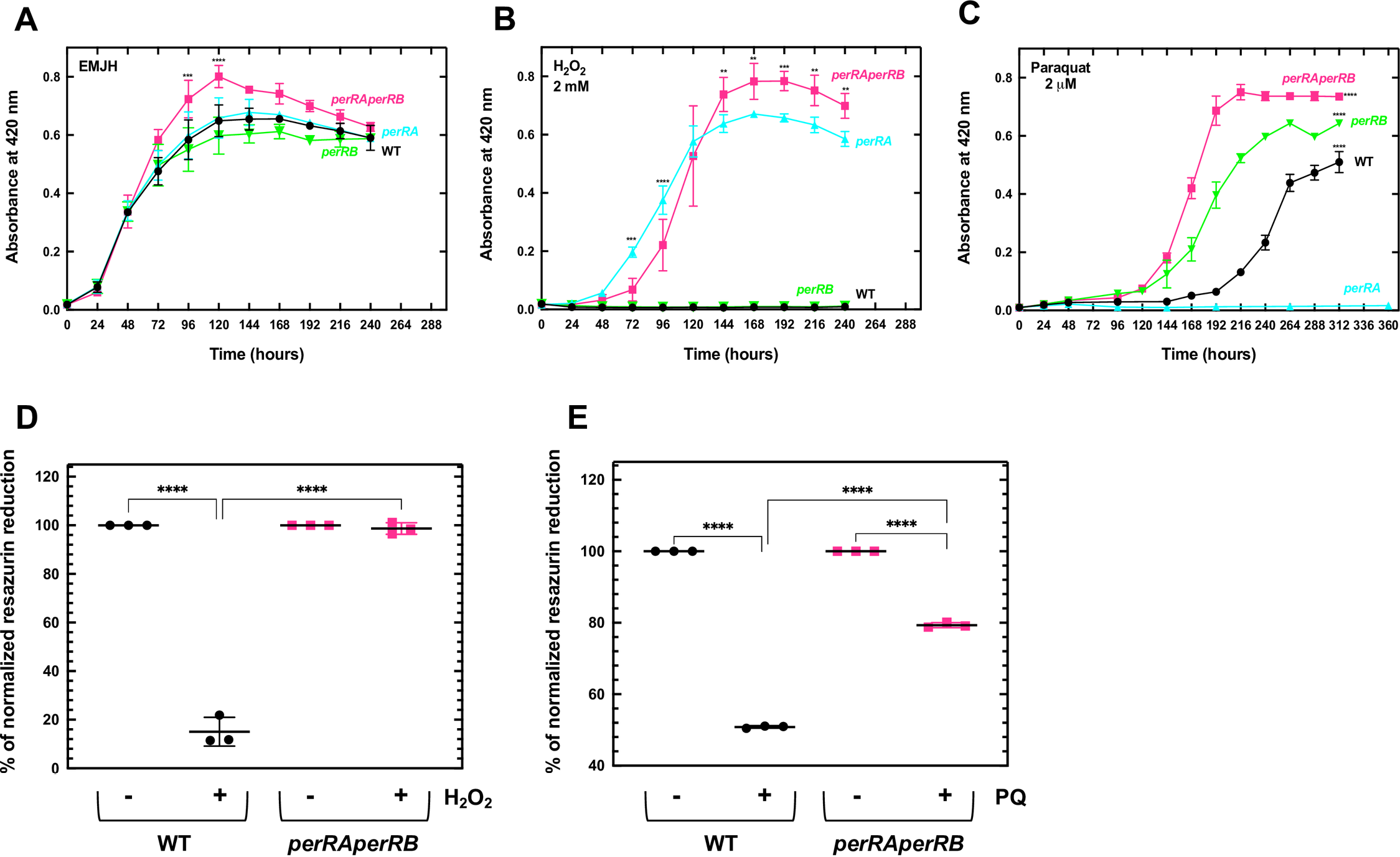
Effect of concomitant inactivation of *perRA* and *perRB* on *Leptospira* growth in the presence of ROS. *L. interrogans* WT (black circles), the single *perRA* mutant (cyan triangles), the single *perRB* mutants (green inverted triangles) or the double *perRAperRB* mutant (pink squares) were cultivated in EMJH medium in the absence (A), or in the presence of 2 mM H2O2 (B) or 2 µM paraquat (C). Growth was assessed by measure of absorbance at 420 nm and the data are means and standard errors of three independent biological replicates. In panel A, statistical significance between the *perRAperRB* mutant and the other strains was determined by Two-way Anova test (***, p-value<0.0008; ****, p-value<0.0001). In panel B, statistical significance between the *perRA* and the *perRAperRB* mutants was determined by two-way Anova test (**, p-value<0.008; ***, p-value<0.0005; ****, p-value<0.0001). In panel C, two-way Anova test indicated that differences between WT, *perRB* mutant, and *perRAperRB* mutant were significant from 144 h (****, p-value<0.0001). Quantitative survival tests were performed by incubating exponentially growing WT (black circles) and double *perRAperRB* mutant (pink squares) with 5 mM H2O2 for 30 min (D) or 100 μM paraquat for 60 min (E). Resazurin reduction was determined as described in the Material and Method section. Data are means and standard errors of three independent biological replicates. Statistical significance was determined by a One-way Anova test (****, p-value<0.0001).

We then tested whether *perRA* and *perRB* inactivation had any influence on *L. interrogans* virulence in the animal model of acute leptospirosis. All hamsters infected intraperitoneally with 10^4^ bacteria of the *perRA* or *perRB* single mutant strains exhibited morbidity sign after 7-8 days, similarly to the WT strain (Fig 7A). In contrast, all animals infected intraperitoneally with 10^4^ or 10^6^ bacteria of the double *perRAperRB* mutant strain did not show any sign of morbidity three weeks post-infection (Fig 7A-B). Moreover, the double *perRAperRB* mutant could not be recovered from kidney and liver of infected animals (Fig 7C-D). Therefore, the double *perRAperRB* mutant exhibited a dramatically reduced virulence and an inability to colonize a host, despite a higher resistance to ROS.

**Fig 7.**
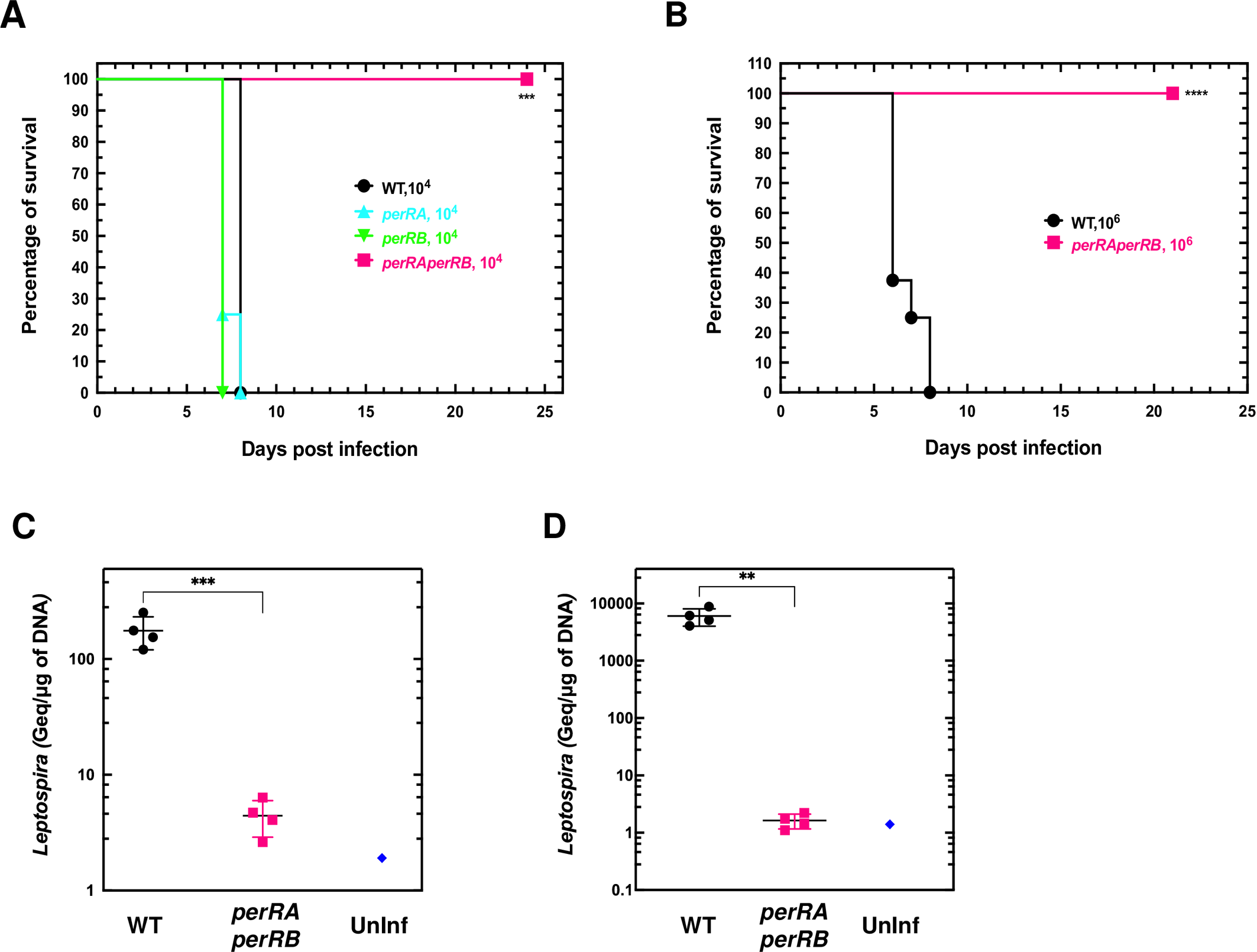
Effect of concomitant inactivation of *perRA* and *perRB* on *Leptospira* virulence. Virulence. (A) was assessed by infecting hamsters (n=4-8) by peritoneal route with 10^4^ (A) or 10^6^ (B) WT (black circles), 10^4^ single *perRA* or *perRB* mutants (cyan triangles and green inverted triangles, respectively), or 10^4^ (A) or 10^6^ (B) double *perRAperRB* mutant (pink squares) strains as described in Material and Methods section. Leptospiral load in kidney (C) and liver (D) of hamsters (n=4) infected with the WT (black circles) or with the *perRAperRB* mutant (pink squares) was assessed by quantitative PCR as described in the Material and Methods section. A sample of non-infected hamsters (blue diamonds) was included as a control. Statistical significance in comparison with WT samples was determined by a Log rank Mantel Cox test (in A, *** p-value=0.0009; in B, **** p-value<0.0001) and by an unpaired t- test (in C, *** p-value=0.0009; in D, ** p-value=0.001).

In spite of several attempts, we could not complement the loss of virulence of the double *perRAperRB* mutant. This prompted us to check the presence of spontaneous mutations in this mutant. Whole DNA sequencing was performed on the WT, *perRB* and double *perRAperRB* mutant strains. As seen in S6 Fig, 2 mutations were found in the double *perRAperRB* mutant that were not present in the *perRB* mutant, its parental strain. One nucleotide insertion was present in the LIMLP_11570 ORF (encoding a 3-oxoacyl ACP synthase putatively involved in fatty acid synthesis) in the *perRAperRB* mutant, however a similar insertion was also observed in several *L. interrogans* isolates from human and animals. One non-synonymous SNP was identified specifically in the double *perRAperRB* mutant. This SNP was located in the LIMLP_01895 ORF, which encodes a hybrid histidine kinase. This missense mutation led to the conservative substitution of Ala with Val at position 148, between the CheY-like response regulator and PAS domains (see in S6 Fig). The resulting protein still displays intact Asp56 and His294, the two putative phosphorylation sites necessary for signal transduction leading to regulation of effectors.

Concomitant inactivation of *perRA* and *perRB* correlates with a pleiotropic effect in *L. interrogans* gene expression.

To further understand the interplay between PerRA and PerRB in controlling the oxidative stress response and virulence in *L. interrogans*, we performed RNA-Seq experiments on the double *perRAperRB* mutant and compared its transcriptomic profile with that of WT and single *perRA* and *perRB* mutant strains.

949 and 1024 ORFs were down- and up-regulated, respectively, in the double *perRAperRB* mutant (Fig 8A and see S3 Table for a complete set of data), which corresponds to almost half of the total coding sequences of *L. interrogans*. In comparison, only about 1% and 3% of the total coding sequences of *L. interrogans* were differentially-expressed in the single *perRA* and *perRB* mutants, respectively (S2 Table) [19]. Volcano scatter plot representation indicated not only a higher magnitude of fold changes but also a greater statistical significance in the double *perRAperRB* mutant (Fig 8B-D).

**Fig 8.**
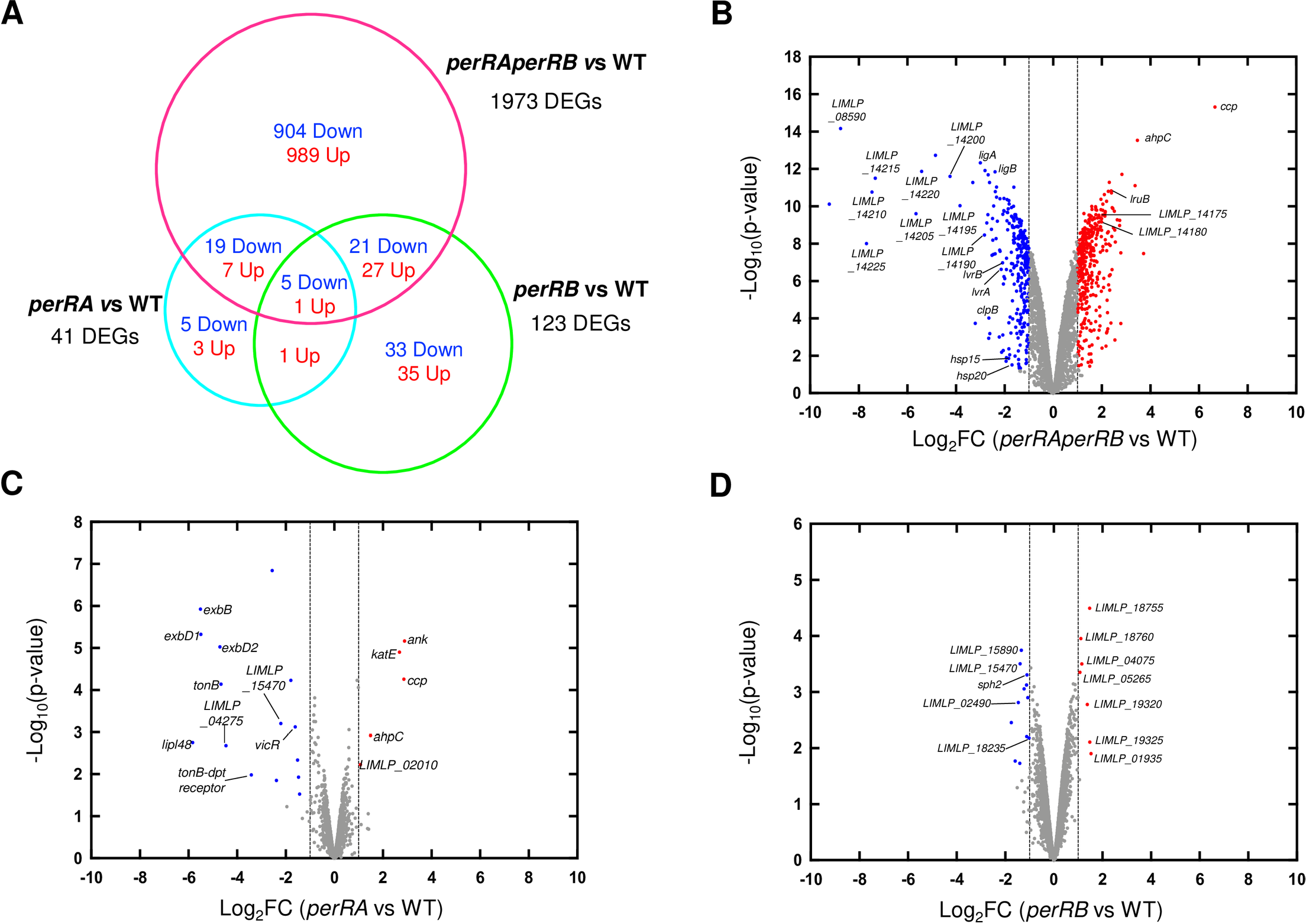
Differential gene expression in the *perRAperRB* mutant. (A) Venn diagram showing the overlap of differentially-expressed ORFs (with an adjusted p-value < 0.05) in the double *perRAperRB* mutant (in pink) with those of the *perRA* mutant (as determined by Zavala-Alvarado *et al.* [19]) (in cyan) and of the *perRB* mutant (as determined in this study) (in green). (B)-(D) Volcano scatter representation of differentially-expressed genes in the *perRAperRB* mutant (B), in the single *perRA* mutant (as determined by Zavala-Alvarado *et al.* [19]) (C), and in the single *perRB* mutant (as determined in this study) (D). Red and blue dots indicate significantly up- and down-regulated genes, respectively, with a Log2FC cutoff of ±1 (dashed vertical lines) and p-value<0.05. Selected genes are labelled.

Most of the differentially-expressed ORFs in the *perRA* mutant were also differentially-expressed in the double *perRAperRB* mutant (Fig 8A). Many genes of the LIMLP_04240- 04285 cluster encoding a putative TonB-dependent transport system, the two-component system VicKR (LIMLP_16720-16725), a putative hemolysin (LIMLP_15470) and several ORFs of unknown function (from LIMLP_14190 to LIMLP_14225) were down-regulated in the *perRA* [12, 19] and *perRAperRB* mutants (Figs 8 and 9A, S4 Table). Likewise, the ORFs encoding the catalase, AhpC and CCP (LIMLP_10145, LIMLP_05955 and LIMLP_02795, respectively), that are repressed by PerRA and up-regulated in the single *perRA* mutant [12, 19], were also up-regulated in the double *perRAperRB* mutant (Figs 8 and 9B, S5 Table).

**Fig 9.**
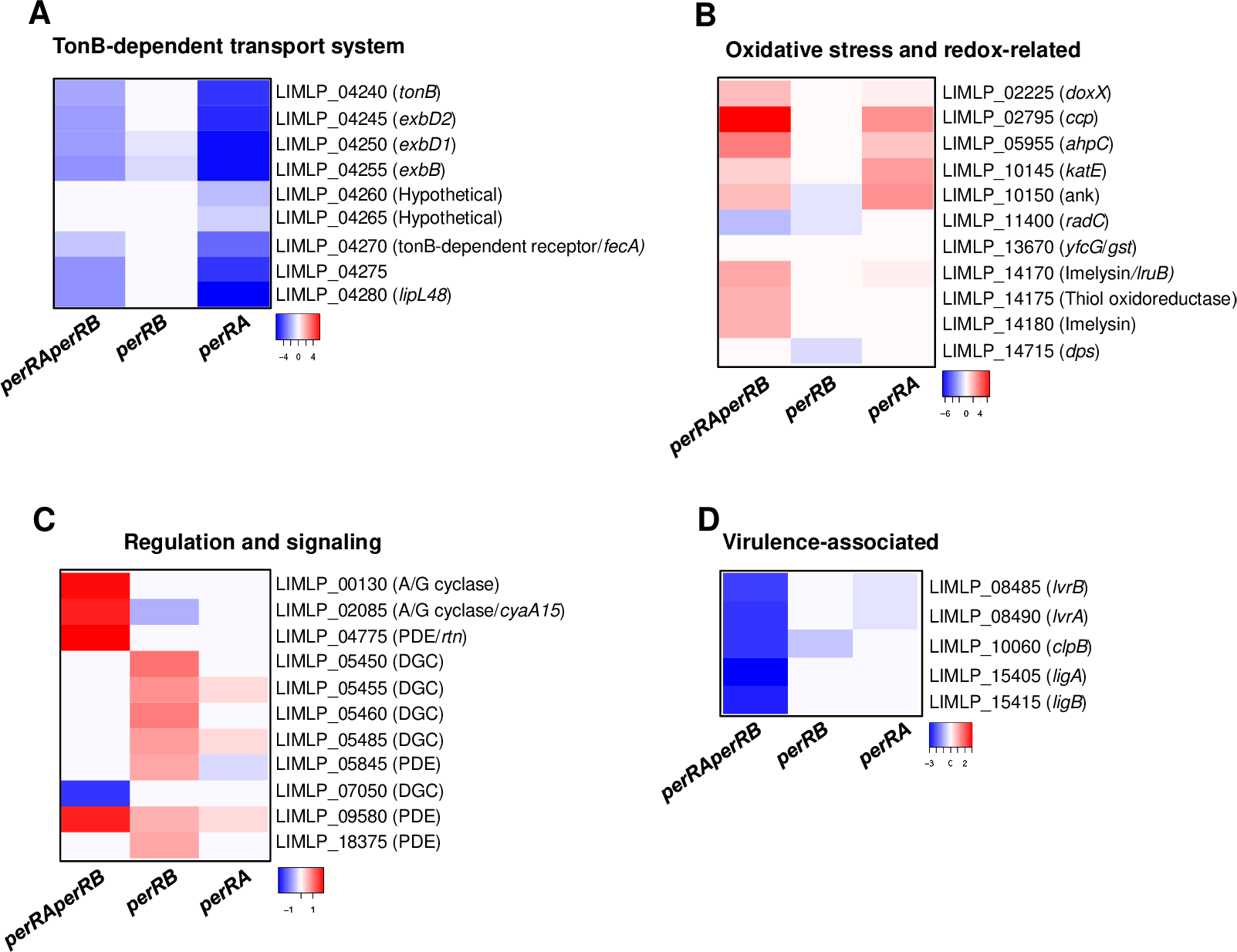
Comparison of differential gene expression in the double *perRAperRB* mutant with that in the single *perRA* and *perRB* mutants. Expression of selected genes of the TonB-dependent transport cluster (A), involved in oxidative stress and redox homeostasis (B), in regulation and signaling (C), and in virulence (D) determined by RNASeq in the double *perRAperRB* mutant was compared to those in the single *perRA* mutant determined by Zavala-Alvarado *et al.* [19] and single *perRB* mutant (as determined in this study). Differential expression in each mutant strain was normalized with that in the WT strain. Gene names are indicated on the right. The Heat Map color from blue to red indicates low to high Log2FC.

21 and 27 ORFs that are respectively down- and up-regulated in the *perRB* mutant were also down- and up-regulated in the *perRAperRB* mutant (Fig 8A). LIMLP_11400 (encoding RadC), LIMLP_04255, encoding ExbB of the TonB-dependent transport system, the hemolysin-encoding ORF LIMLP_15470, and LIMLP_15890 were down-regulated in the single *perRB* and double *perRAperRB* mutants (Fig 8 and S2 and S4 Tables).

Interestingly, the vast majority of the differentially-expressed ORFs in the double *perRAperRB* mutant did not exhibit any change in their expression in the single *perRA* and *perRB* mutants. For instance, among the DGCs and PDEs that were up-regulated in the *perRB* mutant, only LIMLP_09580 was up-regulated in the double *perRAperRB* mutant (S2, S4 and S5 Tables). In fact, the *perRAperRB* double mutant exhibited a distinct expression pattern of genes involved in signaling (Fig 9C). The LIMLP_07050 ORF that codes for a DGC was down-regulated; two ORFs encoding adenylate/guanylate cyclases (LIMLP_00130 and LIMLP_02085) and the PDE-encoding LIMLP_04775 ORF were up-regulated in the *perRAperRB* mutant. Finally, only 6 ORFs were differentially expressed in all mutants (Fig 8A), including LIMLP_04255 and LIMLP_15470 (S4 and S5 Tables). Moreover, a substantial number of regulatory factors (transcriptional regulators, two-component systems, sigma factors) were differentially-expressed exclusively in the *perRAperRB* mutant (S4 and S5 Tables).

In the double *perRAperRB* mutant, several ORFs encoding factors putatively involved in cell growth (cell division, respiration and cell wall homeostasis), chemotaxis and motility are significantly up-regulated, with a Log2FC greater than 1.5 (S5 Table).

In addition to the PerRA-repressed peroxidases (catalase, AhpC, CCP), other oxidative-stress related factors exhibited a higher expression in the *perRAperRB* mutant (Fig 9B). DoxX- encoding ORF, which is up-regulated upon concomitant *perRA* and *perRB* inactivation (Log2FC 1.65), is an integral membrane protein that interacts with SodA in *M. tuberculosis* and participates in tolerance to oxidative and redox stress [31]. Two imelysins (LIMLP_14170/LruB and LIML_14180) and a thiol peroxidase (LIML_14175) exhibited also a higher expression in the *perRAperRB* mutant (Log2FC of 2.36, 1.93, and 2.04 respectively, Fig 9B). All these up-regulated factors (except DoxX) were also up-regulated upon exposure to deadly H2O2 dose [19] and they probably participate in the higher tolerance of the double mutant in the presence of oxidants. Despite the up-regulation of several factors involved in the defense against ROS, the DNA repair protein RadC (encoded by LIMLP_11400) and the glutathione S transferase (encoded by LIMLP_13670) were notably down-regulated in the *perRAperRB* mutant (Log2FC of -1.9 and -2.3, respectively) (Fig 9B and S4 Table).

Strikingly, several down-regulated ORFs in the double *perRAperRB* mutant such as *clpB*, *ligA*, *ligB*, and the operon *lvrAB* have been associated with *Leptospira* virulence. As in many bacteria, leptospiral ClpB ATPase is involved in disaggregating protein aggregates arising upon stress-induced protein denaturation [32]. *ClpB* expression is increased upon exposure to H2O2 and it is required for *Leptospira* virulence [19, 33]. The ClpB-encoding ORF (LIMLP_10060) is dramatically down-regulated in the *perRAperRB* mutant (Log2FC of -2.99) (Fig 8B, Fig 9D and S4 Table).

Another virulence factors in *Leptospira* are the immunoglobulin-like LigA (LIMLP_15405) and LigB (LIMLP_15415) proteins. These surface-exposed proteins are involved in adhesion to host cells through ECM binding [34] and participate in the immune evasion through binding to the host complement Factor H and C4b binding protein [35]. Simultaneous down- regulation of *ligA* and *ligB* expression led to attenuation of *Leptospira* virulence [36]. *LigA* and *ligB* were down-regulated in the *perRAperRB* mutant (Log2FC of -3 and -2.44, respectively) (Fig 8B, Fig 9D and S4 Table).

*LvrA* (LIMLP_08490) and *lvrB* (LIMLP_08485) encode a hybrid histidine kinase and a hybrid response regulator, respectively. Inactivation of the *lvrAB* operon led to virulence attenuation in *L. interrogans* [37]. *LvrA* and *lvrB* had both a decreased expression in the *perRAperRB* mutant (Log2FC of -2.3) (Fig 8B, Fig 9D and S4 Table).

Additional ORFs encoding chaperones (the small heat shock proteins Hsp15 and Hsp20) or enzymes involved in protein folding (the disulfide isomerase DsbD and the peptidyl-prolyl cis-trans isomerase SlyD) and degradation (HtpX) were down-regulated in the *perRAperRB* mutant. DsbD and these two small Hsps were up-regulated in *L. interrogans* upon exposure to H2O2 [19]. All these factors might protect *Leptospira* proteostasis under adverse conditions as encountered during infection inside a host.

The down-regulation of the virulence-associated genes together with the differential expression of several other genes was confirmed by RT-qPCR (S7 Fig).

Taken together, these findings indicate that the concomitant inactivation of *perRA* and *perRB* correlated with global changes in gene expression, leading to the deregulation of several virulence-associated genes.

### Identification of differentially-expressed non-coding RNAs in the *perRB* and *perRAperRB* mutants

Intergenic regions were also analyzed to identify differentially expressed predicted non-coding RNAs (ncRNAs). As observed for coding sequences, inactivation of *perRB* led to the deregulation of only a few putative ncRNAs and most of the changes in expression were below two folds (see S6 Table for a complete set of data). Nonetheless, a few numbers of ncRNAs were significantly down-regulated with a Log2FC below -1 (S7 Table). Some of the differentially-expressed ncRNAs (LepncRNA36, LepncRNA87, LepncRNA89, LepncRNA109, LepncRNA139) were located in the proximate vicinity of differentially-expressed ORFs in the *perRB* mutant. Three ncRNAs (LepncRNA35, LepncRNA89 and LepncRNA109) were also differentially expressed upon *perRA* inactivation (S7 Table) [19].

55 putative ncRNAs were differentially-expressed (with a Log2FC cutoff of ±1) in the *perRAperRB* mutant and several of them were adjacent or overlapped differentially-expressed ORFs (S8 Table). Only a few of these differentially-expressed ncRNAs had an altered expression in the single *perRA* and *perRB* mutant (S8 Table) [19].

Among the most highly differentially-expressed ncRNAs was LepncRNA38 that was located downstream *ccp*, a highly up-regulated ORF in the *perRAperRB* mutant (Fig 10 and S8 Table). LepncRNA38 and *ccp* were also up-regulated in the *perRA* mutant [19]. The ncRNA LepncRNA49, which was down-regulated in the *perRAperRB* mutant, overlapped with *exbB* (LIMLP_04255), an ORF that was also down-regulated in the double *perRAperRB* mutant as well as in the single *perRA* and *perRB* mutants (Fig 10). The down-regulated LepncRNA105 and LepncRNA130 ncRNAs were located downstream the *hsp20-15* operon and *gst*, respectively, three ORFs whose expression is decreased is the *perRAperRB* mutant (Fig 10 and S8 Table). It is worth noting that LepncRNA38, LepncRNA105 and LepncRNA130 are up-regulated by H2O2 as were *ccp*, *hsp20-15* and *gst* ([19]; Fig 10).

**Fig 10.**
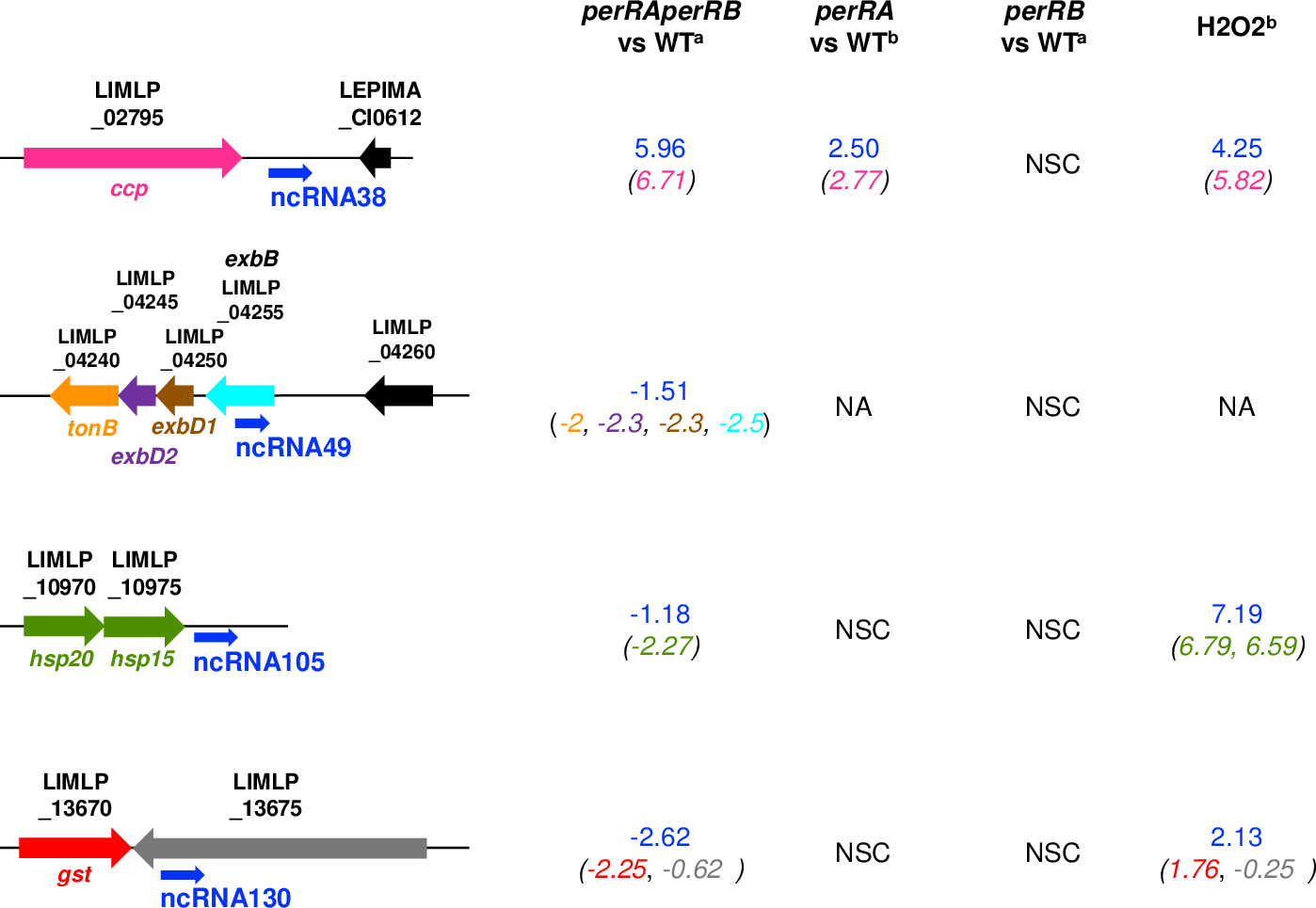
Non-coding RNAs expression in the double *perRAperRB* mutant. Differential expression of selected ncRNAs (LepncRNA38, 49, 105, and 130) in the *perRAperRB* mutant (determined in this study) (^a^) was compared to those in the single *perRA* mutant, as determined by Zavala-Alvarado *et al.* [19] (^b^), in the single *perRB* mutant (determined by this study) (^a^), and upon exposure to 1 mM H2O2 for 1h00 as determined by Zavala-Alvarado *et al.* [19] (^b^). The location of the ncRNAs LepncRNA38, 49, 105, and 130 were represented schematically with the adjacent or overlapping ORFs. The values indicate the Log2FC of ncRNAs expression normalized with that in WT. The respective expression of these ORFs (Log2FC) are indicated into parenthesis with the same color as their representation in the cartoon. NSC, non-significantly changed; NA, non applicable.

Altogether, these findings indicate that the absence of both PerRA and PerRB correlates with major changes in the transcriptional activity of many ncRNAs in *L. interrogans*, that could consequently alter the expression of many ORFs.

## Discussion

Virulence mechanisms are poorly characterized in pathogenic *Leptospira*. These bacteria possess a high number of genes encoding proteins of unknown function (40% of the genomes) and many of them are pathogen-specific. Pathogenic *Leptospira* spp. lack many classical virulence factors, such as type III to type VI secretion systems, and it is unclear which factors are important for their pathogenesis. It is therefore generally agreed that these pathogens possess unique virulence factors. Nonetheless, studying heme oxygenase and catalase mutants have shown that, *in vivo*, iron acquisition and defense against peroxide stress are important virulence-associated mechanisms in *L. interrogans* [11, 38]. Catalase is repressed by PerRA [12, 19] and genes encoding factors involved in iron uptake are very likely controlled by regulators of the Fur-family.

In addition to PerRA, pathogenic *Leptospira* contain three other ORFs annotated as Furs. In the present study, we have characterized the *L. interrogans* Fur-like regulator encoded by LIMLP_05620 and showed that it exhibits characteristic features of a PerR regulator. We consequently named this ORF *perRB*. Sequence alignment and phylogenetic analyses revealed that PerRB is the closest relative to the already characterized PerRA, and perhaps more importantly, they both do exhibit the canonical amino acid residues that are the hallmark of a PerR. The H2O2 sensing histidine and aspartate residues are conserved in *Leptospira* PerRA and PerRB and, interestingly, both genes are H2O2-responsive, albeit with different apparent sensitivity. This is consistent with a mechanism whereby PerRA and PerRB would repress their own transcription and dissociate from their promoter upon oxidation by H2O2, leading to alleviation of repression. Moreover, the higher survival of the *perRB* mutant in the presence of superoxide suggests a derepression of genes encoding defenses against ROS and therefore the participation of PerRB in controlling the adaptation to oxidative stress. Neither *perRA* nor *perRB* expression was up-regulated in iron-limiting condition [12]. Although the putative lipoprotein LIMLP_18755 was significantly up-regulated in the *perRB* mutant and under iron-limiting condition, there was no strong overlap between PerRB regulon and the transcriptional response to iron-limiting condition [12]. Altogether, these findings suggest that LIMLP_05620 encodes a PerR-like rather than a Fur regulator. However, because iron homeostasis and oxidative stress are intertwined, a certain functional relationship has been observed between PerR and Fur. In several bacteria where PerR and Fur coexist, including *B. subtilis* and *C. jejuni*, the PerR regulon overlaps with that of Fur [39, 40]. In addition, *fur* and several Fur-regulated genes are also differentially expressed in the presence of H2O2 [41, 42]. In fact, PerR represses *fur*, whose expression is up-regulated in the presence of H2O2 as a consequence of dissociation of PerR from the *fur* promoter [43, 44]. Metal-catalyzed oxidation of the H2O2 sensing residues will be fundamental in establishing that the PerRB regulator is a *bona fide* PerR.

The coexistence of several PerR regulators in a bacterium is rare. Three PerR-like regulators have been reported only in Gram-positive bacteria such as *B. licheniformis* and *M. smegmatis*. It was shown that the three *B. licheniformis* PerRs sense hydrogen peroxide by histidine oxidation, although with different sensitivity [45]. In *M. smegmatis*, three Fur-like paralogs displayed the canonical PerR Asp residue involved in H2O2 sensitivity, exhibited H2O2 sensing by metal-catalyzed histidine oxidation and a higher H2O2 resistance was observed when their genes were inactivated [46].

One important question was to understand the mechanism that had led to the coexistence of PerRA and PerRB exclusively in highly virulent species (P1 clade). Virulent mammalian- adapted strains in the *Leptospira* genus might have originated from a free-living ancestor inhabiting soils. The phylogenetic analysis presented here indicates that the coexistence of PerRA and PerRB is not due to gene duplication. Indeed, PerRA was already present in the leptospirales ancestor whereas PerRB was probably acquired by the most recent common ancestor of the P1 clade by horizontal transfer from a soil/aquatic bacterium of another phylum (the closest homologues being found in the proteobacteria *Silvanigrella aquatica*). In this scenario, PerRA would had been lost by the P2 clade intermediate species but maintained together with PerRB by the P1 clade species to establish full virulence.

We had previously shown that when *perRA* was inactivated, *L. interrogans* acquired a higher resistance to H2O2 explained by the derepression of *katE*, *ahpC* and *ccp* [12, 19]. Here, we have demonstrated that inactivating *perRB* resulted in a higher survival of *L. interrogans* in the presence of superoxide but it did not affect the survival of *L. interrogans* in the presence of H2O2. Therefore, even though *perRB* is up-regulated upon exposure to H2O2 as *perRA*, the consequence of *perRB* inactivation is different than that of *perRA*, suggesting that PerRA and PerRB have a distinct and non-redundant function in *Leptospira* adaptation to oxidative stress. The distinct distribution of PerRA and PerRB in the *Leptospira* genus and differences in their respective regulon support the hypothesis of a non-redundant function in the adaptation to oxidative stress.

Phenotypic studies suggest that PerRB represses (directly or indirectly) genes encoding defenses against superoxide. Pathogenic *Leptospira* species do not encode any SOD or SOR that could be responsible for detoxification of superoxide whereas saprophyte non-pathogenic *Leptospira* do have such enzymes. Overall, the differentially-expressed genes upon *perRB* inactivation are mostly *Leptospira*-specific and poorly characterized. The highest differentially-expressed ORFs were mainly involved in regulation and cell signaling (transcription and sigma factors, adenylate/diguanylate cyclase, TCSs) and could be involved in regulating the adaptation to various challenging stress encountered within a host. Further studies will be required to determine whether superoxide detoxification in *L. interrogans* is mediated by enzymatic detoxification or metal-dependent scavenging mechanisms and to clarify the exact role of PerRB in controlling those pathways.

The low number of significantly differentially-expressed genes in the *perRB* mutant when *L. interrogans* are cultivated *in vitro* led us to propose that PerRB exerts its function during oxidative stress or upon host-related conditions. Consistent with this hypothesis is the up- regulation of *perRB* in the presence of lethal H2O2 dose. It is worth noting that there is, to some extent, an overlap between the PerRB regulon and the differentially-expressed genes upon exposure to lethal H2O2 dose [19]. Moreover, the exclusive presence of PerRB in the pathogenic *Leptospira* clades strongly suggests that PerRB function is more related to regulating adaptation to infection-related conditions rather than to environmental survival. Consistent with this hypothesis is our observation that the viability of the *perRB* mutant in spring water was comparable to that of the WT (S8 Fig).

One feature of *Leptospira* genus is the genetic and functional redundancy where multiple genes commonly encode for a similar function. The development of genetic tools has made random and targeted mutagenesis possible in *Leptospira*, albeit with a low efficiency in comparison to model bacteria. Due to this limitation, only a few *Leptospira* virulence factors have been identified. The present study is the first to report the concomitant inactivation of two genes. Obtaining a double *perRAperRB* mutant gave us the unique opportunity to investigate the functional relationship between two PerR-like regulators in a pathogenic bacterium.

In many pathogens which contain only one PerR paralog, such as *N. gonorrhoeae*, *S. pyogenes,* and *S. aureus*, PerR was shown to be involved in virulence [44,47–50]. The single *L. interrogans perRA* and *perRB* mutants still retain full virulence in the model for acute leptospirosis (this study and [12]). Interestingly, virulence attenuation was only observed in the double *perRAperRB* mutant, suggesting an interplay in controlling (directly or indirectly) *L. interrogans* virulence-associated genes. We could not successfully complement the double *perRAperRB* mutant. Nevertheless, whole-genome sequencing identified one non- synonymous SNP present only in the double mutant. This SNP resulted in a conservative alanine to valine substitution (both aliphatic amino acids) in the LIMLP_01895 (encoding a hybrid histidine kinase), that should not trigger a drastic functional change in the protein.

Further experiments, including inactivation of the LIMLP_01895-encoded kinase, will be necessary to determine whether it plays any role in *Leptospira* virulence.

The loss of virulence of the double *perRAperRB* mutant correlated with a large differential gene and ncRNA expression compared not only with the WT but also with the single mutant strains. In other words, the double *perRAperRB* displayed differential gene expression that were not observed in the single *perRA* and *perRB* mutants. While we cannot fully exclude a participation of the LIMLP_01895-encoded kinase in some of the differential gene expression, this could indicate that a subset of genes and ncRNAs is controlled by both PerRA and PerRB. The absence of one regulator could be compensated by the other and most of the genes and ncRNAs that can be regulated by the two regulators would not be differentially expressed in the single mutants. The few genes (LIMLP_02010, LIMLP_04255, LIMLP_04235, LIMLP_11810, LIMLP_14225, LILP_15470, and LIMLP_18235) and ncRNAs that display differential expression in the single mutants in our transcriptomic studies (this study and [19]) indicate a certain functional redundancy of the two regulators even if the phenotypic analyses of the mutants suggest distinct functions. The change in expression of a few regulators when PerRA and PerRB are both absent could lead to major changes in expression of many ORFs or ncRNAs in cascade. The differential expression of many regulators in the double mutant could also explain that the transcriptional profile of the double mutant is not simply the cumulative gene deregulation of the respective single *perRA* and *perRB* mutants.

Despite a higher ability to resist ROS, the double *perRAperRB* mutant lost its virulence; it could not trigger acute leptospirosis-associated morbidity. This could be obviously explained by a significant lower expression of several virulence-associated factors in the double *perRAperRB* mutant, such as LigA, LigB, LvrA, LvrB, and ClpB. In addition, other dramatically down-regulated genes encode factors such as small Hsps (Hsp15 and Hsp20) for which a role in bacterial virulence is demonstrated in other bacteria including *M. tuberculosis* [51, 52]. Moreover, several differentially-expressed ORFs of unknown function could also be responsible for the loss of virulence of the double *perRAperRB* mutant.

In summary, this study has allowed to identify a second PerR-like regulator in pathogenic *L. interrogans* strains that cooperates with PerRA to control the adaptation to oxidative stress. By concomitantly inactivating *perRA* and *perRB*, we have unveiled a complex regulatory network that could reveal a putative functional relationship between PerR regulators and *Leptospira* virulence, most likely through the regulation of virulence- and pathogenicity- associated factors.

## Materials and Methods

### Bacterial strains and growth condition

*L. interrogans* serovar Manilae strain L495, *perRA* (M776), *perRB* (M1474), and the double *perRAperRB* mutant strains (see S9 Table for a complete description of the strains used in this study) were grown aerobically at 30°C in Ellinghausen-McCullough-Johnson-Harris medium (EMJH) [53] with shaking at 100 rpm. *Leptospira* growth was followed by measuring the absorbance at 420 nm. *β*2163 and *Π*1 *E. coli* strains were cultivated at 37°C in Luria-Bertani medium with shaking at 37°C in the presence of 0.3 mM thymidine or diaminopimelic acid (Sigma-Aldrich), respectively. When needed, spectinomycin and kanamycin were added at the respective concentration of 50 and 30 µg/ml.

### Determination of bacteria viability

Leptospires survival was determined by incubating exponentially growing *L. interrogans* cells (≈ 10^8^/ml) in EMJH in the presence or absence of 100 μM paraquat for 60 min or 5 mM H2O2 for 30 min at 30°C. Colony-forming unit quantification was performed by diluting bacteria in EMJH and plating on solid EMJH medium. After one month incubation at 30°C, colonies were counted and percent survival (% of colony forming unit, CFU) was calculated as the ratio of CFU for bacteria incubated in the presence of ROS to that for bacteria incubated in the absence of ROS. For the colorimetric assay, 100 µl of leptospires were incubated with resazurin (Alamar Blue reagent, ThermoScientific) for 48h. Leptospires viability was assessed by their capacity to reduce blue resazurin into pink resorufin as described previously [54].

### Concomitant inactivation of *perRA* (LIMLP_10155) and *perRB* (LIMLP_05620)

*PerRA* gene (LIMLP_10155/LIC12034/LA1857) was inactivated in the *perRB* (LIMLP_05620/LIC11158/LA2887) mutant strain (M1474, *perRB::Km^R^*) by introduction of a spectinomycin resistance cassette (S5 Fig). For this, a spectinomycin resistance cassette flanked by 0.8 kb sequences homologous to the sequences flanking *perRA* was created by gene synthesis (GeneArt, Life Technologies) and cloned into a kanamycin-resistant *Escherichia coli* vector unable to replicate in *Leptospira*. The obtained suicide plasmid (pK*Δ*perRA) (S10 Table) was introduced in the *perRB* mutant strain by electroporation as previously described [55] using a Gene Pulser Xcell (Biorad). Individual spectinomycin-resistant colonies were selected on EMJH plates containing 50 µg/ml spectinomycin and screened by PCR (using the P1 and P2 primer set, see S11 Table) for proper replacement of the *perRA* coding sequence by the spectinomycin resistance cassette. *PerRA* inactivation in the double *perRAperRB* mutant was verified by western blot using an anti-PerRA serum (S5Fig).

### Complementation of the *perRB* mutant

The *perRB* mutant *(perRB::Km^R^*) complementation was performed by expressing the *perRB* ORF in the pMaORI replicative vector [56]. The *perRB* (LIMLP_05620) ORF together with its native promoter region (200 bp upstream region) were amplified from genomic DNA of *L. interrogans* serovar Manilae strain L495 (using the ComPerR2_5Not and ComPerR2_3Xba primer set, S11 Table) and cloned between the Not1 and Xba1 restriction sites in the pMaORI vector. The absence of mutation in the *perRB* locus in the obtained plasmid (pNB139) was checked by DNA sequencing and the pNB139 plasmid was introduced in the *perRB* mutant (M1474) by conjugation using the *E. coli* β2163 conjugating strain as previously described [57]. *Leptospira* conjugants were selected on EMJH plates containing 50 µg/ml spectinomycin and resistant colonies were then inoculated into liquid EMJH medium supplemented with spectinomycin for further analysis. The restoration of PerRB production in the complemented *perRB* mutant was verified by western blot (S3 Fig).

### Phylogenetic analyses

The sequences homologous to the LIMLP_10155 (PerRA), LIMLP_05620 (PerRB), LIMLP_18590 and LIMLP_04825 proteins were searched with BLASTP version 2.10.0 among the other *Leptospira* species (Fig 3 and S1 Table) or among the protein sequences of 11,070 representative genomes (Fig 2), as previously described [58]. In that case, only the sequences with an e-value less than 1e-10 and a percentage of similarity greater than 60% were retained. Sequences with percent identity equal to 100% were clustered by CD-HIT version 4.8.1 and only one sequence was retained. The resulting 1671 sequences were subsequently aligned by MAFFT version 7.471. A phylogenetic tree was finally built with IQ- TREE version 2.1.1 under the best-fit model LG + R10. A second phylogenetic tree was made with a subset of sequences to improve the resolution of the separation between PerRA and PerRB. The same procedure was followed, except that the best-fit model used for phylogenetic reconstruction is LG + R5. Both trees were visualized with FigTree version 1.4.4 (https://github.com/rambaut/figtree).

### RNA purification

Virulent *L. interrogans* serovar Manilae strain L495 and *perRB* (M1474) mutant strains with less than three *in vitro* passages were used in this study. Four independent biological replicates of exponentially grown *L. interrogans* WT, *perRB* (M1474) and double *perRAperRB* mutant strains were harvested and resuspended in 1 ml TRIzol (ThermoFisher Scientific) and stored at -80°C. Nucleic Acids were extracted with chloroform and precipitated with isopropanol as described earlier [59]. Contaminating genomic DNA was removed by DNAse treatment using the Turbo DNA-free kit (ThermoFisher Scientific) as described by the manufacturer. The integrity of RNAs (RIN > 8.0) was verified by the Agilent Bioanalyzer RNA NanoChips (Agilent technologies, Wilmington, DE).

### RNA Sequencing

rRNA were depleted from 0.5 µg of total RNA using the Ribo-Zero rRNA Removal Kit (Bacteria) from Illumina. Sequencing libraries were constructed using the TruSeq Stranded mRNA Sample preparation kit (20020595) following the manufacturer’s instructions (Illumina). The directional libraries were controlled on Bioanalyzer DNA1000 Chips (Agilent Technologies) and concentrations measured with the Qubit dsDNA HS Assay Kit (ThermoFisher). Sequences of 65 bases were generated on the Illumina Hiseq 2500 sequencer.

Reads were cleaned of adapter sequences and low-quality sequences using cutadapt version 1.11 [60]. Only sequences at least 25 nt in length were considered for further analysis. Bowtie version 1.2.2 [61], with default parameters, was used for alignment on the reference genome (*L. interrogans* serovar Manilae strain UP-MMC-NIID LP, from MicroScope Platform). Genes were counted using featureCounts version 1.4.6-p3 [62] from Subreads package (parameters: -t gene -g locus_tag -s 1).

Count data were analyzed using R version 3.5.1 [63] and the Bioconductor package DESeq2 version 1.20.0 [64]. The normalization and dispersion estimation were performed with DESeq2 using the default parameters and statistical tests for differential expression were performed applying the independent filtering algorithm. Differential expressions were expressed as logarithm to base 2 of fold change (Log2FC). A generalized linear model including the replicate effect as blocking factor was set in order to test for the differential expression between *Leptospira* samples. Raw p-values were adjusted for multiple testing according to the Benjamini and Hochberg (BH) procedure [65] and genes with an adjusted p-value lower than 0.05 and a Log2FC higher than 1 or lower than -1 were considered differentially expressed. Heat maps and Volcano plots were generated using the Galaxy platform (https://usegalaxy.eu/).

### Quantitative RT-PCR experiments

cDNA synthesis was performed with the cDNA synthesis kit (Biorad) according to the manufacturer’s recommendation. Quantitative PCR was conducted in triplicate with the SsoFast EvaGreen Supermix (Biorad) as previously described [19]. *FlaB* (LIMLP_09410) was chosen as reference gene.

### Non-coding RNA identification

Sequencing data from the *L. interrogans* WT, *perRB* (M1474) and double *perRAperRB* mutant strains were processed with Trimmomatic [66] to remove low-quality bases and adapter contaminations. BWA mem (version 0.7.12) was used to discard the reads matching *Leptospira* rRNA, tRNA or polyA sequences and to assign the resulting reads to *Leptospira* replicons. Then Rockhopper [67] was used to re-align reads corresponding to separate replicons and to assemble transcripts models. The output was filtered to retain all transcripts longer than 50 nucleotides not overlapping within 10 nucleotides with NCBI annotated genes on the same orientation, and showing a minimum Rockhopper raw count value of 50 in at least two isolates. This high-quality set of new sRNA was subjected to differential expression analysis with Rockhopper, adopting a Benjamini-Hochberg adjusted P-value threshold of 0.01. For each non-coding RNAs, putative function was identified by BLAST using the Rfam database [68].

### Infection experiments

WT and mutant *L. interrogans* strains were cultivated in EMJH medium until the exponential phase and counted under a dark-field microscope using a Petroff-Hauser cell. 10^4^ or 10^6^ bacteria (in 0.5 ml) were injected intraperitoneally in groups of 4-8 male 4 weeks-old Syrian Golden hamsters (RjHan:AURA, Janvier Labs). Animals were monitored daily and sacrificed by carbon dioxide inhalation when endpoint criteria were met (sign of distress, morbidity).

To assess leptospiral load in tissues, liver and kidneys were sampled from infected hamsters (n=4) (with 10^6^ leptospires) 6-8 or 21 days post-infection for the WT and the *perRAperRB* mutant, respectively. Genomic DNA was extracted from about 35 mg of tissue using the Maxwell™ 16 Tissue DNA purification kit (Promega) and total DNA concentrations were normalized to 20 ng/µl. Leptospiral load was assessed by quantitative PCR using the SsoFast EvaGreen Supermix (Biorad) with *lipL32* primers (S11 Table). The PCR reactions were run with the CFX96™ Real-Time System (Biorad) using the absolute quantification program as follows: 95°C for 3 min followed by 40 cycles of amplification (95°C for 10 s and 55°C for 30 s). Genome equivalents (Geq) were calculated as previously described [69].

### Ethics Statement

The protocol for animal experimentation was reviewed by the Institut Pasteur (Paris, France), the competent authority, for compliance with the French and European regulations on Animal Welfare and with Public Health Service recommendations. This project has been reviewed and approved (CETEA #2016-0019) by the Institut Pasteur ethic committee for animal experimentation, agreed by the French Ministry of Agriculture.

## Acknowledgement

The authors would like to thank Melissa Caimano and André Grassmann for fruitful discussions.

## Funding

Crispin Zavala-Alvarado and Samuel Garcia Huete were supported by the Pasteur - Paris University (PPU) International PhD Program. This project has received funding from the European Union’s Horizon 2020 research and innovation programme (https://ec.europa.eu/programmes/horizon2020/en) under the Marie Sklodowska-Curie grant agreement No 665807 (https://ec.europa.eu/research/mariecurieactions). Crispin Zavala-Alvarado has received funding from Fondation Etchebès-Fondation France (S-CM 16008) (https://www.fondationdefrance.org/fr/fondation/fondation-etchebes).

The funders had no role in study design, data collection and analysis, decision to publish, or preparation of the manuscript.

## Supporting information

**S1 Fig. Phylogenetic analysis of the four Fur-like regulators of *L. interrogans*.**

Extended phylogenetic tree showing the separation between PerRA (LIMLP_10155 in red) and PerRB (LIMLP_05620 in blue).

**S2 Fig. Growth of *L. interrogans perRA* and *perRB* mutants in the presence of H2O2.**

*L. interrogans* WT (black circles), *perRA* (cyan triangles) and *perRB* (green inverted triangles) mutant strains were cultivated in EMJH medium at 30°C in the absence (A) or presence of 2 mM H2O2 (B). *Leptospira* growth was assessed by absorbance at 420 nm. Data are means and standard errors of three independent biological experiments.

**S3 Fig. Complementation of the *perRB* mutant.**

*L. interrogans* WT, *perRB* mutant and *perRB* mutant complemented strains were cultivated in EMJH medium at 30°C until the logarithmic phase and lyzed by sonication in 25 mM Tris pH 7.5, 100 mM KCl, 2 mM EDTA, 5 mM DTT, with a protease inhibitors cocktail (cOmplete Mini EDTA-free, Roche). 10 μg of total lyzates were resolved on a 15% SDS-PAGE and transferred on nitrocellulose membrane. PerRB was detected by immunoblot (A) using a rabbit polyclonal antibody at 1/500 dilution. FcpA production was assessed as a control of equal loading using a rabbit polyclonal antibody at 1/1000 dilution. A goat anti-rabbit IgG secondary antibody coupled to the HRP peroxidase (Sigma) was used at a 1/150000 dilution. Detection was performed by chemiluminescence with the Supersignal^TM^ West Pico PLUS reagent (ThermoScientific). PerRB content was quantified using ImageJ (Schneider *et al*., Nat Methods **9,** 671–675 (2012) https://doi.org/10.1038/nmeth.2089) and normalized by the quantity in the WT strain (B). Data are means and standard errors of three independent biological experiments. Statistical significance was determined by a One-way Anova test in comparison with WT samples (**, p-value=0.0018).

**S4 Fig. Analysis of the *L. interrogans* PerRB regulon.**

(A) Venn diagram showing the overlap of differentially-expressed ORFs (with an adjusted p-value < 0.05) in the *perRA* and *perRB* mutants. Differentially-expressed genes in the *perRB* mutant (as determined in this study) (in green) were compared with those in the *perRA* mutant as determined previously (Zavala-Alvarado *et al*., PLOS Pathogens. 2020 Oct 6;16(10):e1008904) (in cyan). The down- and up-regulated ORFs in both mutants were indicated in blue and red, respectively. (B) Comparison of differentially-expressed ORFs (with an adjusted p-value < 0.05) in the *perRB* mutant and upon *L. interrogans* exposure to H2O2. Log2FC of differentially-expressed ORFs upon *L. interrogans* exposure to 1 mM H2O2 (as determined previously (Zavala-Alvarado *et al*., PLOS Pathogens. 2020 Oct 6;16(10):e1008904)) was plotted against the Log2FC of differentially-expressed ORFs upon *perRB* inactivation. Down- and up-regulated ORFs in the *perRB* mutant were represented by blue and red symbols, respectively, and the name of selected ORFs was indicated. The dashed lines indicate a Log2FC value of zero. Please note that only differentially-expressed ORFs in both conditions were considered.

**S5 Fig. Concomitant inactivation of *perRA* and *perRB*.**

(A) Schematic representation of the double *perRAperRB* mutant construction. *PerRA* (LIMLP_10155) was inactivated by allelic exchange in the transposon *perRB* mutant. The kanamycin (Km) and spectinomycin (Spc) resistance cassettes inactivating *perRB* and *perRA*, respectively, are indicated.

(B) Production of PerRA in the WT, in the single *perRA* and *perRB* mutants and in the double *perRAperRB* mutant strains. *L. interrogans* strains were cultivated in EMJH medium at 30°C until the logarithmic phase and lyzed by sonication in 25 mM Tris pH 7.5, 100 mM KCl, 2 mM EDTA, 5 mM DTT, with a protease inhibitors cocktail (cOmplete Mini EDTA-free, Roche). 10 μg of total lyzates were resolved on a 15% SDS-PAGE and transferred on nitrocellulose membrane. PerRA was detected by immunoblot using a 1/2000 antibody dilution as described previously (Kebouchi *et al*., J Biol Chem. 2018;293(2):497-509. doi:10.1074/jbc.M117.804443).

(C) Growth of stationary phase-adapted WT and *perRAperRB* mutant strains. *L. interrogans* WT (black circles) and *perRAperRB* mutant (pink squares) were cultivated in EMJH medium at 30°C until late stationary phase (7 days after the entry in the stationary phase) and used to inoculate EMJH medium. Bacteria were then cultivated at 30°C and growth was assessed by absorbance at 420 nm. Data are means and standard errors of three independent biological experiments.

**S6 Fig. Mutations found in the *perRAperRB* mutant compared to the parental strain (*perRB* mutant).**

(A) Genomic DNA of *perRB* mutant and *perRAperRB* mutant strains was extracted with the Maxwell™ 16 cell DNA purification kit (Promega) and sequenced by Next-generation sequencing. Sequence Reads were processed by fqCleaner and aligned with the reference sequenced genome of *Leptospira interrogans* serovar Manilae strain UP-MMC-NIID LP (accession numbers CP011931, CP011932, CP011933; Satou *et al.*, Genome Announc. 2015 Aug 13;3(4):e00882-15. doi: 10.1128/genomeA.00882-15. PMID: 26272567; PMCID: PMC4536678.) using Burrows-Wheeler Alignment tool (BWA mem 0.7.5a) (Li & Durbin, Bioinformatics. 2009;25(14):1754-1760. doi:10.1093/bioinformatics/btp324). SNP and Indel calling was done with the Genome Analysis Tool Kit GATK2 following the Broad Institute best practices (McKenna *et al.*, Genome Res. 2010;20(9):1297-1303. doi:10.1101/gr.107524.110) The mutation identified (indicated in red) were further confirmed by sequencing PCR-amplified DNA fragments. 358 *L. interrogans* genomes of a cgMLST *Leptospira* isolates database (https://bigsdb.pasteur.fr/leptospira/) (Guglielmini *et al.*, PLoS neglected tropical diseases, (2019) *13*(4), e0007374. https://doi.org/10.1371/journal.pntd.0007374) were screened for homolog of the affected ORFs using BLASTN 2.9.0+ (Altschul *et al.*, Nucleic Acids Res. 1997;25(17):3389-3402. doi:10.1093/nar/25.17.3389). Identified mutations were searched by alignment using MAFFT version 7.453 (Katoh & Standley, Molecular Biology and Evolution, Volume 30, Issue 4, April 2013, Pages 772–780, https://doi.org/10.1093/molbev/mst010. *^a^* Positions, reference sequences, and annotations were indicated according to the *L. interrogans* serovar Manilae strain UP-MMC-NIID LP (MicroScope Platform (https://mage.genoscope.cns.fr/microscope/home/index.php). *^b^* Isolates Id 38, 747, 806, 816, 974, 986, 1058, 1085.

(B) Schematic representation of LIMLP_01895.

Schematic representation and domain organization of LIMLP_01895 have been determined by ScanProsite (De Castro *et al*., Nucleic Acids Res. 2006 Jul 1;34(Web Server issue):W362- 5.) Nucleotide and corresponding amino acid sequences were indicated from residues 145 to 151 with the mutated codon and amino acid in red. The substituted nucleotide was underlined.

**S7 Fig. RT-qPCR experiments in the double *perRAperRB* mutant.**

RNAs were extracted from exponentially-grown *L. interrogans*strains WT or double *perRAperRB* mutant (*m*). Expression of the indicated genes was measured by RT-qPCR using the LIMLP_06735 as reference gene. This gene, which encodes a protein with unknown function, was used as a reference gene since its expression was not changed upon inactivation of *perRA* and *perRB*. Gene expression in the *perRAperRB*mutant was normalized against that in the WT strain. Fold change values are indicated in blue. Statistical significance was determined by a Two-way Anova test in comparison with the WT samples (****, p- value<0.0001; **, p-value=0.0048).

**S8 Fig. Survival of *perRA* and *perRB* mutants in spring water.**

Exponentially growing WT (black circles), *perRA* (cyan triangles) and *perRB*(green squares) mutant strains were centrifugated at 2600 g for 15 min and washed three times and resuspended into filter-sterilized spring water (Volvic). All samples were adjusted to 5x10^8^ leptospires/ml. The samples were incubated at RT in darkness and, at the indicated times, leptospires were counted under a dark-field microscope using a Petroff-Hauser cell (A) and their viability was determined by quantification of resazurin reduction using the AlamarBlue reagent (ThermoScientific) (B). Data are means and standard errors of three independent biological experiments. Statistical significance was determined by a Two-way Anova test in comparison with the WT samples (**, p-value=0.006; *, p-value=0.0278).

S1 Table. Distribution of the four Fur-like regulators of *Leptospira interrogans* in the genus *Leptospira*.

S2 Table. Complete set of ORF expression in *Leptospira interrogans* WT and M1474 *perRB* mutant.

S3 Table. Complete set of ORF expression in *Leptospira interrogans* WT and double *perRAperRB* mutant.

S4 Table. Selected down-regulated genes in the *perRAperRB* double mutant.

S5 Table. Selected up-regulated genes in the *perRAperRB* double mutant.

S6 Table. Complete set of differentially-expressed predicted non coding RNAs in the *perRB* (M1474) and in the double *perRAperRB* mutant strains of *Leptospira interrogans*.

S7 Table. Selected differentially-expressed non-coding RNAs in the *perRB* mutant

S8 Table. Selected differentially-expressed non-coding RNAs in the *perRAperRB* mutant.

S9 Table. Strains used in this study

S10 Table. Plasmids used in this study

S11 Table. Primers used in this study

## Notes

### Competing Interest Statement

The authors have declared no competing interest.

