## Supplementary material for "The oxidative stress response of pathogenic *Leptospira* is controlled by two peroxide stress regulators which putatively cooperate in controlling virulence": S4Table_ZavalaAlvarado_revised_D-20-0288.docx

| **ORF ID^a^** | **ORF**  ***L. biflexa*^b^** | **Gene** | **Function** | **Log_2_FC** | **Adjusted**  **p-value** | **Log2FC**  ***perRA* vs WT^c^** | **Log2FC**  ***perRB* vs WT^d^** |
| --- | --- | --- | --- | --- | --- | --- | --- |
| ***Oxidative stress and redox-related*** |  |  |  |  |  |  |  |
| LIMLP_11400 (LIC12297/LA1456)* |  |  | DNA repair protein RadC | -1.905 | 9.64e-53 |  | -0.619 |
| LIMLP_13670 (LIC10807-LEPIC0823/LA3356)* |  | *gst/*  *yfcG* | Glutathione S transferase | -2.251 | 3.88e-37 |  |  |
| ***Regulation and signaling*** |  |  |  |  |  |  |  |
| LIMLP_02535 (LIC12980/LA0598) | *LEPBI_I0893* |  | Transcriptional regulator | -2.339 | 1.85e-54 | 0.586 |  |
| LIMLP_07050 (LIC11444/LA2528)* |  | *pleD* | Diguanylate cyclase | -1.511 | 1.18e-37 |  |  |
| LIMLP_08485 (LIC11708/LA2223)* |  | *lvrB* | Two component System Histidine kinase LvrB | -2.305 | 1.92e-26 |  |  |
| LIMLP_08490 (LIC11709/LA2222)* |  | *lvrA* | Two component System Response Regulator LvrA | -2.377 | 1.68e-26 |  |  |
| LIMLP_11545 (LIC12319/LA1428)* |  |  | Serine Threonine Phosphatase | -2.697 | 5.06e-108 |  |  |
| LIMLP_12520 (LIC12505/LA1185)* |  |  | Response regulator | -1.597 | 1.84e-42 |  |  |
| LIMLP_16720 (LIC13269/LA4102) | LEPBI_I3226 | *vicR* | Response regulator | -2.238 | 6.90e-55 | -1.611 |  |
| LIMLP_16725 (LIC13270/LA4104) | LEPBI_I3227 | *vicK* | Signal transduction histidine kinase | -1.493 | 1.27e-19 | -0.919 |  |
| ***Chaperones, protein folding and quality control*** |  |  |  |  |  |  |  |
| LIMLP_08595 (LIC11731/LA2194) | LEPBI_I1734 | *slyD* | Peptidyl-prolyl cis-trans isomerase | -1.804 | 15.05e-46 |  |  |
| LIMLP_10060 (LIC12017/LA1879)* | *LEPBI_I2449* | *clpB* | ClpB molecular chaperone | -2.996 | 2.21e-25 |  |  |
| LIMLP_10970 (LIC12210/LA1564)* | LEPBI_I1849 | *ibpa* | Small Heat Shock Protein Hsp20 | -2.273 | 2.73e-02 |  |  |
| LIMLP_10975 (LIC12215/LA1563)* | LEPBI_I1848 | *hsp15* | Small Heat Shock Protein Hsp15 | -2.274 | 9.76e-03 |  |  |
| LIMLP_11965 (LIC12404/LA1321)* | *LEPBI_I3297* | *dsbD* | Thiol disulfide interchange protein | -1.896 | 4.12e-72 |  |  |
| LIMLP_18560 (LIC20141/LB174)* | LEPBI_II0128 | *htpX* | Protease HtpX | -1.609 | 2.22e-51 |  |  |
| ***Miscellaneous*** |  |  |  |  |  |  |  |
| LIMLP_01290 (LIC13232/LA4052) |  | *dltE* | Short-chain dehydrogenase | -1.592 | 1.16e-38 |  |  |
| LIMLP_01575 (LIC13177/LA3974)* | *LEPBI_I2304* |  | Exonuclease | -1.651 | 3.38e-15 |  |  |
| LIMLP_02840 (LIC12921/LA0678)* | LEPBI_I2436 | *mcpA* | Methyl-accepting chemotaxis protein | -3.183 | 1.10e-85 |  |  |
| LIMLP_03360 (LIC12818/LA0802)* |  | *pilF* | TPR-containing protein/pilus assembly | -1.721 | 1.28e-50 |  |  |
| LIMLP_04240 (LIC10889/LA3247) | LEPBI_I0146 | *tonB* | Energy transporter TonB | -2.062 | 2.91e-77 | -4.601 |  |
| LIMLP_04245 (LIC10890/LA3246)* | LEPBI_I0147 | *exbD2* | Biopolymer transport protein ExbD/TolR | -2.284 | 4.51e-66 | -4.606 |  |
| LIMLP_04250 (LIC10891/LA3245) | LEPBI_I0148 | *exbD1* | Biopolymer transport protein ExbD/TolR | -2.309 | 8.65e-40 | -5.355 |  |
| LIMLP_04255 (LIC10892/LA3244) | LEPBI_I0149 | *exbB* | Biopolymer transport protein ExbB/TolQ | -2.554 | 2.82e-74 | -5.478 | -0.947 |
| LIMLP_08585 (LEPIC1765-LIC11730/LA2196) |  |  | Stage II sporulation protein E | -1.975 | 1.73e-16 |  |  |
| LIMLP_08600 (LIC11732/LA2193)** |  | *pcrA/*  *uvrD* | ATP-dependent DNA helicase | -1.620 | 3.54e-49 |  |  |
| LIMLP_09815 (LA1937)* |  |  | Transposase | -2.269 | 3.49e-22 |  |  |
| LIMLP_12135 (LIC12436/LA1276) |  |  | Sulfatase | -2.571 | 1.11e-42 |  |  |
| LIMLP_14200 (LIC10707/LA3474)* |  |  | GDSL-like lipase | -4.169 | 1.23e-05 | -0.853 |  |
| LIMLP_15000 (LIC10548/LA3668)* |  |  | Uracil-DNA glycosylase A | -1.678 | 3.15e-47 |  |  |
| LIMLP_15405 (LIC10465/LA3778)* |  | *ligA* | Immunglobulin-like LigA | -3.005 | 6.32e-172 |  |  |
| LIMLP_15415 (LIC10464/LA3778)* |  | *ligB* | Immunglobulin-like LigB | -2.444 | 3.21e-111 |  |  |
| LIMLP_15470 (LIC10454/LA3793)* | LEPBI_I0671 |  | Putative hemolysin | -1.865 | 2.53e-64 | -2.154 | -1.317 |
| LIMLP_16555 (LIC10239/LA0283)* | *LEPBI_I2991* |  | Beta propeller repeat protein | -2.201 | 3.71e-77 |  |  |
| LIMLP_16870 (LIC13298/LA4137)* | *LEPBI_I0609* |  | NADPH-dependent FMN reductase | -1.715 | 2.05e-35 |  |  |
| LIMLP_18070 (LIC20049/LB063)* | LEPBI_II0035 |  | NAD kinase | -2.318 | 5.54e-102 |  |  |
| LIMLP_18085 (LIC20052/LB068)* |  | *ole* | Fatty acid desaturase | -1.700 | 7.27e-16 |  |  |
| LIMLP_19560 (LA1821) |  | *cysE* | Serine acetyl transferase | -1.573 | 5.60e-11 |  |  |
| ***Hypothetical*** |  |  |  |  |  |  |  |
| LIMLP_00370 (LIC10068/LA0075)* |  |  | Hypothetical | -2.929 | 1.82e-101 |  |  |
| LIMLP_02040 (LIC13081/LA3856)* |  |  | Hypothetical | -1.988 | 8.61e-11 |  |  |
| LIMLP_02135 (LIC13058/LA0495)** | *LEPBI_I3000* |  | Hypothetical | -1.949 | 8.99e-18 |  |  |
| LIMLP_03245 (LA0774)* |  |  | Hypothetical | -2.435 | 8.38e-29 |  |  |
| LIMLP_03485 (LIC12794/LA0829) |  |  | Hypothetical | -1.784 | 1.41e-32 |  |  |
| LIMLP_03490 (LIC12794/LA0829) |  |  | Hypothetical | -1.678 | 4.25e-50 |  |  |
| LIMLP_04015 (LIC12692/LA0958)* | LEPBI_I1783 |  | Hypothetical | -1.749 | 3.35e-49 |  |  |
| LIMLP_04260 (LIC10893/LA3243) |  |  | Hypothetical | -1.690 | 8.90e-02 | -1.519 |  |
| LIMLP_04275 (LIC10897/LA3241) |  |  | Hypothetical | -2.505 | 2.91e-25 | -3.888 |  |
| LIMLP_04280 (LIC10898/LA3240) |  | *lipl48* | Hypothetical lipoprotein | -2.572 | 3.73e-81 | -5.506 |  |
| LIMLP_04285 (LIC10899/LA3239) |  |  | Hypothetical | -2.687 | 4.45e-43 |  |  |
| LIMLP_04465* |  |  | Hypothetical | -1.632 | 2.19e-04 |  |  |
| LIMLP_04480* |  |  | Hypothetical | -2.383 | 4.38e-44 |  |  |
| LIMLP_04510 (LA3180)* |  |  | Hypothetical | -1.967 | 3.93e-06 |  |  |
| LIMLP_05215 (LIC11078/LA2896)* |  |  | Hypothetical | -1.971 | 5.85e-08 |  |  |
| LIMLP_05995 (LIC11228/LA2796)* |  |  | Hypothetical | -1.640 | 3.09e-28 |  |  |
| LIMLP_06320 (LEPIC1323/LA2720)* |  |  | Hypothetical | -2.114 | 2.59e-55 |  |  |
| LIMLP_06325 (LIC11289/LA2719)* |  |  | Hypothetical putative membrane protein | -1.611 | 9.11e-46 |  |  |
| LIMLP_07040 (LIC11442/LA2530)* |  |  | Hypothetical putative exported protein | -1.673 | 2.86e-15 |  |  |
| LIMLP_08235 |  |  | Hypothetical | -1.512 | 2.84e-03 |  |  |
| LIMLP_08590 (LEPIC1767/LA2195)* |  |  | Hypothetical | -8.924 | 0.00e-01 | -0.859 |  |
| LIMLP_09650 (LIC11935/LA1968) |  |  | Hypothetical outer membrane protein | -2.186 | 1.26e-37 | -1.787 |  |
| LIMLP_09995 (LIC12005/LA1894)* |  |  | Hypothetical | -2.261 | 2.74e-24 |  |  |
| LIMLP_10015* |  |  | Hypothetical | -1.571 | 1.07e-05 |  |  |
| LIMLP_10255 (LIC12073/LA1730)* | LEPBI_I2295 |  | Hypothetical | -1.545 | 3.04e-12 |  |  |
| LIMLP_11405 (LIC12298/LA1455)* | LEPBI_I0934 |  | Hypothetical | -1.935 | 1.51e-29 |  | -0.577 |
| LIMLP_11805 (LIC12370) |  |  | Hypothetical | -1.704 | 2.25e-02 |  |  |
| LIMLP_11810 (LIC12371/LA1359) |  |  | Hypothetical | -1.469 | 3.19e-34 | -0.760 | -0.600 |
| LIMLP_13620 (LIC10816/LA3344)* |  |  | Hypothetical putative membrane protein | -5.008 | 4.17e-229 |  |  |
| LIMLP_14190 (LIC10709/LEPIN3051)** |  |  | Hypothetical lipoprotein | -2.632 | 1.88e-05 | -0.679 |  |
| LIMLP_14195 (LIC10708/LA3473)** |  |  | Hypothetical | -3.717 | 1.32e-12 | -0.813 |  |
| LIMLP_14205 (LIC10706/LA3475)** |  |  | Hypothetical lipoprotein | -5.967 | 4.75e-02 | -0.847 |  |
| LIMLP_14210 (LIC10705/LA3477)** |  |  | Hypothetical lipoprotein | -7.278 | 3.05e-03 | -0.915 |  |
| LIMLP_14215 (LIC10704/LA3478)** |  |  | Hypothetical putative lipoprotein | -7.343 | 3.05e-03 |  |  |
| LIMLP_14220 (LIC10703/LA3479) |  |  | Hypothetical | -5.317 | 1.13e-02 |  |  |
| LIMLP_14225 (LIC10702/LA3480)* |  |  | Hypothetical | -7.918 | 2.14e-03 | -0.775 | -0.879 |
| LIMLP_14905 (LIC10567/LA3642) |  |  | Hypothetical | -1.573 | 2.95e-02 |  |  |
| LIMLP_15420 (LIC10463/LA3779) |  |  | Hypothetical putative lipoprotein | -1.640 | 2.42e-31 |  |  |
| LIMLP_15425 (LIC10462/LA3780) |  |  | Hypothetical putative lipoprotein | -1.882 | 2.32e-16 |  |  |
| LIMLP_15430 (LIC10461/LA3781)** |  |  | Hypothetical putative lipoprotein | -2.375 | 3.48e-99 |  |  |
| LIMLP_15890 (LIC10377/LA0430) |  |  | Hypothetical putative exported protein | -1.954 | 3.70e-37 |  | -1.353 |
| LIMLP_16575 (LIC10235/LA0278)* | LEPBI_I3216 |  | Hypothetical | -1.818 | 4.84e-43 |  |  |
| LIMLP_16765 (LEPIC3342/LEPIN3585)* |  |  | Hypothetical | -1.800 | 9.62e-38 |  |  |
| LIMLP_17480 (LIC13428/LA4282) |  |  | Hypothetical | -1.678 | 3.63e-42 |  |  |
| LIMLP_18235(LIC20078/LB099) |  |  | Hypothetical | -0.559 | 2.91e-07 | -0.658 | -1.098 |
| LIMLP_18725 (LIC20172/LB217)* | LEPBI_II0156 |  | Hypothetical putative lipoprotein | -2.644 | 5.80e-128 |  |  |
| LIMLP_19195 (LIC20259/LB340)* |  |  | Hypothetical | -1.981 | 3.45e-03 |  |  |
| LIMLP_19265 (LB360) |  |  | Hypothetical | -1.787 | 3.34e-45 |  |  |
| LIMLP_19335 (LA1773) |  |  | Hypothetical putative membrane protein | -2.551 | 4.09e-31 |  |  |

**S4 Table. Selected down-regulated genes in the *perRAperRB* double mutant.**

Significantly down-regulated genes upon concomitant inactivation of *perRA* and *perRB*.

^a^ Gene numeration is according to Satou et al. (Satou *et al*., 2015).

^b^ ORF in italic indicates an absence of synteny

^c^ Log_2_FC of significantly differentially-expressed genes (adj. p-value < 0.05) in the *perRA* mutant (M776) (Zavala-Alvarado *et al.*, 2020)

^d^ Log_2_FC of significantly differentially-expressed genes (adj. p-value < 0.05) in the *perRB* mutant (M1474) (this study)

* Up-regulated ORFs upon exposure to 1 mM H_2_O_2_ (adj. p-value < 0.05) (Zavala-Alvarado *et al.*, 2020)

** Down-regulated ORFs upon exposure to 1 mM H_2_O_2_ (adj. p-value < 0.05) (Zavala-Alvarado *et al.*, 2020)
