## Supplementary material for "The oxidative stress response of pathogenic *Leptospira* is controlled by two peroxide stress regulators which putatively cooperate in controlling virulence": S5Table_ZavalaAlvardo_revised_D-20-0288.docx

| **ORF ID^a^** | ***L. biflexa^b^*** | **Gene** | **Function** | **Log_2_FC** | **Adjusted**  **p-value** | **Log2FC**  ***perRA* vs WT^c^** | **Log2FC**  ***perRB* vs WT^d^** |
| --- | --- | --- | --- | --- | --- | --- | --- |
| ***Oxidative stress and redox-related*** |  |  |  |  |  |  |  |
| LIMLP_02225 (LIC13041/LA0520)** | *LEPBI_I1384* |  | DoxX family protein | 1.648 | 2.76e-15 | 0.466 |  |
| LIMLP_02795 (LIC12927/LA0666)* | *LEPBI_I2430* | *ccp* | Cytochrome C peroxidase | 6.712 | 0.00e01 | 2.773 |  |
| LIMLP_05955 (LIC11219/LA2809)* | LEPBI_I1358 | *ahpC* | Peroxiredoxin/alkylperoxiredoxin reductase | 1.539 | 1.23e-05 | 1.539 |  |
| LIMLP_10995 (LIC12215/LA1555)** | LEPBI_I3024 |  | Multicopper oxidase | 1.654 | 3.05e-34 |  |  |
| LIMLP_10145 (LIC12032/LA1859)* |  | *katE* | Catalase | 1.119 | 2.04e-27 | 2.637 |  |
| LIMLP_10150 (LIC12033/LA1858)* |  | *ank* | Ankyrin repeat-containing protein | 1.625 | 3.65e-30 | 2.867 |  |
| LIMLP_14170 (LIC10713/LA3469)* |  | *lruB*  *irpA* | Imelysin (LruB) | 2.356 | 9.02e-95 |  |  |
| LIMLP_14175 (LIC10712/LA3470)* |  |  | Thiol oxidoreductase | 1.930 | 5.20e-44 |  |  |
| LIMLP_14180 (LIC10711/LA3471)* |  |  | Imelysin | 2.043 | 1.54e-43 |  |  |
| ***Regulation/Signaling*** |  |  |  |  |  |  |  |
| LIMLP_00130 (LIC10024/LA0027)** | *LEPBI_I2644* |  | Adenylate/guanylate cyclase | 1.707 | 3.55e-35 |  |  |
| LIMLP_02085 (LIC13072/LA3843)* | *LEPBI_I1085* | *cyaA15* | Adenylate/guanylate cyclase | 1.557 | 1.54e-23 |  |  |
| LIMLP_04775 (LIC10996/LA3104)* |  | *rtn* | Cyclic diguanylate phosphodiesterase (EAL) domain protein | 1.778 | 1.30e-40 |  |  |
| LIMLP_05770 (LIC11191/LA2845) |  |  | Transcriptional regulator | 2.462 | 1.16e-22 |  |  |
| LIMLP_06010 (LIC11231/LA2790)** | *LEPBI_I0100* |  | AcrR family transcriptional regulator | 1.825 | 6.68e-45 |  |  |
| LIMLP_06955 (LIC11425/LA2549)** | *LEPBI_I2119* | *baeS* | GHKL domain protein/  Two-component sensor Histidine kinase | 2.077 | 4.81e-69 |  |  |
| LIMLP_09580 (LIC11921/LA1983) | LEPBI_I1269 | *rtn* | Cyclic diguanylate phosphodiesterase (EAL) domain protein | 1.513 | 3.72e-17 |  | 0.547 |
| LIMLP_10240 (LIC12070/LA1734)** | LEPBI_I2292 |  | Anti-sigma factor antagonist | 2.002 | 1.35e-27 |  |  |
| LIMLP_14970 (LIC10552/LA3662) |  |  | Two component hybrid sensor histidine kinase and regulator | 1.575 | 3.72e-26 |  | 0.972 |
| LIMLP_16165 (LIC10319/LA0372)** | LEPBI_I2371 |  | CopG family transcriptional regulator | 2.294 | 3.51e-07 |  |  |
| LIMLP_18365 (LIC20104/LB130)** |  |  | AraC family transcriptional regulator | 1.546 | 5.06e-07 |  |  |
| LIMLP_19135 (LB325)** | LEPBI_II0039 |  | AcrR family transcriptional regulator | 2.047 | 8.65e-40 |  |  |
| ***Cell division and respiration*** |  |  |  |  |  |  |  |
| LIMLP_03705 (LIC12752/LA0884)** | LEPBI_I1307 | *nuoN* | NADH-quinone oxidoreductase subunit N | 1.704 | 1.34e-27 |  |  |
| LIMLP_03710 (LIC12751/LA0885)** | LEPBI_I1306 | *nuoM* | Proton translocating NADH-quinone oxidoreductase subunit M | 1.806 | 9.18e-56 |  |  |
| LIMLP_05130 (LIC11061/LA3011)** | *LEPBI_I1393* | *ftsK* | DNA translocase FtsK | 1.984 | 2.18e-71 |  |  |
| LIMLP_06175 (LIC11262/LA2754)** | LEPBI_I3076 | *rodA*  *mreA*  *mrdB* | Rod shape-determining protein RodA | 1.514 | 8.36e-26 |  |  |
| LIMLP_09265 (LIC11865/LA2049)** | LEPBI_I1758 | *ftsW* | Cell division protein FtsW | 1.660 | 1.53e-16 |  | 0.645 |
| LIMLP_17445 (LIC13421/LA4275)** | LEPBI_I2421 | *hyfE* | Formate hydrogenase subunit E | 1.808 | 1.28e-20 |  |  |
| LIMLP_17450 (LIC13422/LA4276)** | LEPBI_I2420 | *hyfF* | Formate hydrogenase subunit F/NADH ubiquinone complex I subunit | 1.671 | 6.62e-33 |  |  |
| LIMLP_17460 (LIC13424/LA4278)** | LEPBI_I2417 | *hycG* | hydrogenase 4 subunit G/NADH ubiquinone oxidoreductase | 1.655 | 3.65e-45 |  |  |
| ***Cell wall homeostasis*** |  |  |  |  |  |  |  |
| LIMLP_13245 (LIC12646/LA1009)** | LEPBI_I0637 | *mrcA* | Carboxypeptidase/Transglycosylase/penicillin binding protein | 1.572 | 8.09e-52 |  |  |
| LIMLP_17195 (LIC13365/LA4213)** | LEPBI_I3335 | *nlpD* | M23 family metallo-endopeptidase | 2.000 | 1.25e-59 |  |  |
| LIMLP_17600 (LIC13453/LA4311)** | LEPBI_I3428 |  | M23 family metallo-endopeptidase | 2.061 | 7.55e-21 |  |  |
| LIMLP_19130 (LIC20247/LB323) | LEPBI_II0046 |  | Lytic transglycosylase | 2.416 | 8.03e-45 |  |  |
| ***Motility and chemotaxis*** |  |  |  |  |  |  |  |
| LIMLP_02700 (LIC12947) |  |  | Endoflagellar protein | 1.961 | 2.33e-20 |  |  |
| LIMLP_05655 (LIC11168/LA2876)** | LEPBI_I2922 |  | Chemotaxis phosphatase CheX | 1.868 | 6.03e-41 |  |  |
| LIMLP_06490 (LIC11325/LA2666)** | LEPBI_I1533 | *flgA* | Flagella basal body P-ring formation protein FlgA | 1.705 | 5.65e-40 |  |  |
| LIMLP_06495 (LIC11326/LA2665)** | LEPBI_I1534 | *flgH* | Flagellar L-ring protein FlgH | 1.578 | 5.44e-38 |  |  |
| LIMLP_06500 (LIC11327/LA2664)** | LEPBI_I1535 | *flgI* | Flagellar P-ring protein FlgI | 1.579 | 9.71e-31 |  |  |
| LIMLP_06505 (LIC11328/LA2663)** | LEPBI_I1536 | *flgJ* | Rod binding protein/flagellar protein FlgJ | 1.826 | 6.13e-28 |  |  |
| LIMLP_06865 (LIC11407/LA2574)* | *LEPBI_I2077* |  | Methyl-accepting chemotaxis protein | 2.297 | 2.38e-108 |  |  |
| LIMLP_07475 (LIC11531/LA2418) | LEPBI_I1589 | *flaB* | Flagellar filament 35 kDa core protein | 1.599 | 9.20e-54 |  |  |
| LIMLP_11310 (LIC12278/LA1478)** | LEPBI_I1333 |  | Flagellar assembly protein FlaA | 2.347 | 1.07e-34 |  |  |
| LIMLP_14615 (LIC10624/LA3574)** | LEPBI_I0198 | *fliL* | Flagellar basal body-associated protein FliL | 1.765 | 8.93e-24 |  |  |
| LIMLP_14620 (LIC10624/LA3575)** | LEPBI_I0198 | *fliL* | Flagellar basal body-associated protein FliL | 1.753 | 1.55e-32 |  |  |
| LIMLP_14625 (LIC10623/LA3576)** | LEPBI_I0197 | *motB* | Sodium translocating bacterial flagellar motor protein MotB | 1.730 | 1.40e-58 |  |  |
| LIMLP_14630 (LIC10622/LA3577)** | LEPBI_I0196 | *motA* | Motility protein A | 2.059 | 1.13e-75 |  |  |
| LIMLP_14635 (LIC10621/LA3578)** | LEPBI_I0195 | *flbD* | Flagellar protein FlbD | 1.936 | 6.19e-52 |  |  |
| LIMLP_17580 (LIC13449/LA4309)** | LEPBI_I3211 | *fliW* | Flagellar assembly protein FliW | 1.504 | 8.88e-19 |  |  |
| ***Protein synthesis and secretion*** |  |  |  |  |  |  |  |
| LIMLP_03170 (LIC12855/LA0757)** | LEPBI_I1947 | *rpmD* | 50S ribosomal subunit protein L30 | 1.610 | 1.05e-30 |  |  |
| LIMLP_03220 (LIC12845/LA0766)** | LEPBI_I1938 | *rplQ* | 50S ribosomal subunit protein L17 | 1.595 | 2.12e-17 |  |  |
| LIMLP_07675 (LIC11572/LA2373)** | LEPBI_I1677 | *gspF* | Type II secretion system protein F | 1.570 | 1.33e-30 |  |  |
| LIMLP_12685 (LIC12537/LA1143)** | LEPBI_I1402 |  | Preprotein translocase subunit SecF | 1.697 | 8.17e-11 |  |  |
| ***Miscellaneous*** |  |  |  |  |  |  |  |
| LIMLP_00280 (LIC10053/LA0060)** | LEPBI_I3241 |  | TPR repeat-containing protein | 1.807 | 5.90e-31 |  |  |
| LIMLP_01295 (LIC13231/LA4051)** | *LEPBI_I0067* |  | Sulfatase-modifying factor | 1.735 | 2.62e-33 |  |  |
| LIMLP_01550 (LIC13182/LA3980)* | *LEPBI_I1842* | *ugpQ* | Glycerophosphodiester phosphodiesterase | 2.065 | 3.18e-17 |  |  |
| LIMLP_02820 (LIC12924/LA0673)** | LEPBI_I0184 | *spsF* | Cytidylyltransferase/  spore coat biosynthesis protein F | 1.563 | 4.68e-20 |  |  |
| LIMLP_03875 (LIC12719/LA0927)** | LEPBI_I0867 |  | TPR repeat-containing protein | 1.950 | 2.55e-63 |  |  |
| LIMLP_04740 (LIC10900/LA3112)** |  | *kdpA* | Potassium translocating ATPase subunit A | 1.917 | 4.04e-28 |  |  |
| LIMLP_04965 (LIC11028/LA3067)** | LEPBI_I0909 | *tolB* | Biopolymer transporter TolR | 1.761 | 7.98e-60 |  |  |
| LIMLP_05635 (LIC11164/LA2880)** |  |  | HEPN domain-containing protein/toxin-antitoxin system/DNA-binding protein | 1.579 | 2.76e-07 |  | 0.621 |
| LIMLP_05645 (LIC11166/LA2878) | LEPBI_I2920 | *smtA* | SAM-dependent methyl transferase | 2.203 | 5.31e-53 |  |  |
| LIMLP_06015 (LIC11232/LA2789) | LEPBI_I0789 |  | Aldolase | 1.519 | 5.65e-40 |  |  |
| LIMLP_06385 (LIC11301/LA2703)** | LEPBI_I1799 |  | Sporulation and spore germination | 1.634 | 3.55e-35 |  |  |
| LIMLP_06480 (LIC11323/LA2668)** | *LEPBI_I1344* |  | Methyl transferase domain protein | 2.615 | 2.00e-48 |  |  |
| LIMLP_07110 (LIC11459/LA2509)** | LEPBI_I1732 | *wcaJ* | Sugar transferase | 1.564 | 6.27e-33 |  |  |
| LIMLP_07200 (LIC11479/LA2483) | LEPBI_I2357 | *xerD* | Tyrosine recombinase XerD | 1.865 | 8.61e-27 |  |  |
| LIMLP_08755 (LIC11764/LA2156)** | LEPBI_I2314 |  | Asp amino transferase class I/II | 1.557 | 4.06e-31 |  |  |
| LIMLP_10105 (LIC12024/LA1867) | LEPBI_I1721 |  | TPR repeat-containing protein | 2.113 | 1.29e-43 |  |  |
| LIMLP_10150 (LIC12033/LA1858)* | *LEPBI_I3361* | *ank* | Ankyrin repeat-containing protein | 1.625 | 3.64e-30 | 2.867 |  |
| LIMLP_11840 (LIC12378/LA1350)** | LEPBI_I1267 | *ubiG* | 3-demethylubiquinone-9 3-O-methyltransferase | 2.772 | 1.64e-41 |  |  |
| LIMLP_12045 (LIC12419/LA1299)** | *LEPBI_I3359* |  | Ankyrin repeat-containing protein | 1.538 | 7.13e-28 |  |  |
| LIMLP_12680 (LIC12536/LA1144)** | LEPBI_I1403 | *ribD* | Pyrimidine deaminase zinc-binding region/riboflavin biosynthesis | 1.528 | 1.25e-31 |  |  |
| LIMLP_13345 (LIC10867/LA3275)* |  |  | Toxin-antitoxin system, antitoxin component | 1.921 | 5.83e-11 |  |  |
| LIMLP_13370 (LIC10864/LA3284)** | LEPBI_I2604 |  | TPR repeat-containing protein | 1.992 | 1.38e-38 |  |  |
| LIMLP_13530 (LIC10831/LA3320)** |  |  | LRR protein | 1.749 | 4.93e-12 |  |  |
| LIMLP_13535 (LIC10830/LA3321)** |  |  | LRR protein | 1.913 | 1.09e-10 |  |  |
| LIMLP_14610 (LIC10625/LA3573)** | LEPBI_I0199 | *kdsB* | 3-deoxy-manno-octulosanate cytidylyltransferase | 1.506 | 9.91e-33 |  |  |
| LIMLP_14640 (LIC10620/LA3579)** | LEPBI_I0194 |  | Glycosyl transferase | 2.518 | 1.49e-55 |  |  |
| LIMLP_15050 (LIC10537/LA3685)** | LEPBI_I2664 |  | OmpA | 1.534 | 5.29e-36 |  |  |
| LIMLP_16925 (LIC13309/LA4149)** | LEPBI_I2718 |  | Thioesterase | 1.731 | 5.52e-10 |  |  |
| LIMLP_17200 (LIC13366/LA4215)** | *LEPBI_I2637* |  | Strictosidine synthase | 2.119 | 2.83e-28 |  |  |
| LIMLP_18705 (LIC20168/LB212)** | LEPBI_II0259 |  | Phosphoribosyl ATP-pyrophosphatase | 1.992 | 2.50e-21 |  |  |
| LIMLP_19465 (LA1798) | LEPBI_I0049 |  | Antitoxin component (PHD family) of the type II toxin-antitoxin system | 1.824 | 1.87e-03 |  |  |
| ***Hypothetical*** |  |  |  |  |  |  |  |
| LIMLP_00075 (LIC10015/LA0016) | LEPBI_I0013 |  | Hypothetical | 1.650 | 8.44e-15 |  |  |
| LIMLP_00870 (LIC10166/LA0191) |  |  | Hypothetical | 1.815 | 2.50e-03 |  |  |
| LIMLP_00875 (LEPIC0174/LA0192) |  |  | Hypothetical | 2.035 | 1.94e-06 |  |  |
| LIMLP_00880 (LIC10167/LA0193)* |  |  | Hypothetical | 1.540 | 4.76e-05 |  |  |
| LIMLP_00885 (LEPIC0176/LA0194)* |  |  | Hypothetical | 2.091 | 2.97e-26 |  |  |
| LIMLP_00905 (LIC10170/LA0200) |  |  | Hypothetical | 1.719 | 3.14e-10 |  |  |
| LIMLP_00910 (LIC10171/LA0201) |  |  | Hypothetical | 1.901 | 2.82e-06 |  |  |
| LIMLP_00920 (LIC10173/LA0203) |  |  | Hypothetical | 1.834 | 2.57e-07 |  |  |
| LIMLP_00930 (LIC10175/LA0205)** |  |  | Hypothetical | 1.644 | 3.24e-35 |  |  |
| LIMLP_00935 (LIC10176/LA0206)** |  |  | Hypothetical | 1.679 | 1.78e-16 |  |  |
| LIMLP_00940 (LIC10177/LA0207)** |  |  | Hypothetical | 1.557 | 2.01e-16 |  |  |
| LIMLP_00945 (LEPIC10190/LA0208)** |  |  | Hypothetical | 1.611 | 1.18e-11 |  |  |
| LIMLP_01125 (LIC10213/LA0248)** | LEPBI_I2677 |  | Hypothetical | 1.849 | 1.05e-29 |  |  |
| LIMLP_02145 (LA0497)** |  |  | Hypothetical | 2.174 | 1.11e-03 |  |  |
| LIMLP_02405 (LIC13005/LA0564) |  |  | Hypothetical | 2.782 | 2.44e-45 |  |  |
| LIMLP_02750 (LIC12936/LA0654)** | LEPBI_I3081 |  | Hypothetical | 2.218 | 3.50e-72 |  |  |
| LIMLP_03225 |  |  | Hypothetical | 1.876 | 5.56e-05 |  |  |
| LIMLP_04325 (LIC10906/LA3230) |  |  | Hypothetical putative membrane protein | 0.559 | 1.83e-02 | 0.587 | 0.635 |
| LIMLP_04580 (LIC10957/LEPIN2778) | LEPBI_I1286 |  | Hypothetical | 2.583 | 1.77e-20 |  |  |
| LIMLP_04585 (LA3159) |  |  | Hypothetical | 1.705 | 5.48e-04 |  |  |
| LIMLP_04610 (LIC10963/LA3150)** |  |  | Hypothetical | 1.684 | 3.95e-09 |  |  |
| LIMLP_04620 (LIC10965/LA3148)* |  |  | Hypothetical | 1.785 | 5.98e-05 |  |  |
| LIMLP_04970 (LIC11030/LA3064) |  |  | Hypothetical putative exported lipoprotein | 3.436 | 1.77e-130 |  |  |
| LIMLP_05115 (LIC11059/LA3016)* |  |  | Hypothetical | 2.041 | 5.36e-10 |  |  |
| LIMLP_05120 (LEPIC1091/LA3015)* |  |  | Hypothetical | 1.648 | 1.53e-08 |  |  |
| LIMLP_05650 (LIC11167/LA2877)** |  |  | Putative lipoprotein | 2.144 | 1.25e-20 |  |  |
| LIMLP_07255 (LIC11490/LA2467) | LEPBI_I2658 |  | Hypothetical | 1.717 | 1.53e-25 |  |  |
| LIMLP_07690 (LIC11575/LA2370)** | LEPBI_I1680 |  | Hypothetical | 1.575 | 6.18e-33 |  |  |
| LIMLP_07970 (LIC11631/LA2308)** | LEPBI_I1641 |  | Hypothetical | 2.445 | 3.70e-38 |  |  |
| LIMLP_08065 (LIC11653/LA2285)** | LEPBI_I1467 |  | Hypothetical putative exported protein | 1.623 | 7.90e-32 |  |  |
| LIMLP_08420 (LIC11696/LA2240)* | LEPBI_I2761 |  | Hypothetical | 2.112 | 1.71e-25 |  |  |
| LIMLP_08465 (LEPIC1741/LA2228)** | *LEPBI_I1556* |  | Hypothetical | 2.813 | 8.14e-46 |  |  |
| LIMLP_08470 (LEPIC1742/LEPIN1974) |  |  | Hypothetical | 2.203 | 1.88e-06 |  |  |
| LIMLP_08475 (LIC11706/LA2227)** | LEPBI_I1563 |  | Hypothetical | 2.258 | 2.88e-19 |  |  |
| LIMLP_08760 (LIC11765/LA2155) |  |  | Hypothetical | 2.289 | 1.21e-14 |  |  |
| LIMLP_09380 (LIC11888/LA2020)** |  |  | Hypothetical | 2.897 | 1.24e-127 |  |  |
| LIMLP_09385 |  |  | Hypothetical | 2.457 | 6.59e-07 |  |  |
| LIMLP_09920 (LIC11989/LA1916)** | LEPBI_I2965 |  | Hypothetical putative exported protein | 2.238 | 1.06e-52 |  |  |
| LIMLP_10080 (LIC12020/LA1874)** | LEPBI_I1720 |  | Hypothetical | 1.785 | 4.02e-19 |  |  |
| LIMLP_10245 (LIC12071/LA1733)** | LEPBI_I2293 |  | Hypothetical | 1.775 | 2.27e-38 |  |  |
| LIMLP_11000 (LIC12216/LA1554)** | LEPBI_I3025 |  | Hypothetical putative membrane protein | 1.597 | 2.90e-14 |  |  |
| LIMLP_11230 (LIC12262/LA1496)** | LEPBI_I1030 |  | Hypothetical exported protein | 2.375 | 4.25e-26 |  |  |
| LIMLP_11235 (LIC12263/LA1495)** | LEPBI_I1029 |  | Hypothetical exported protein | 2.319 | 1.42e-65 |  |  |
| LIMLP_11970 (LIC12405/LA1319)** | *LEPBI_I1421* |  | Hypothetical | 1.724 | 3.72e-24 |  |  |
| LIMLP_12545 (LIC12509/LA1180)* |  |  | Hypothetical | 1.891 | 6.77e-10 |  |  |
| LIMLP_13055 (LIC12609/LA1058) |  |  | Hypothetical | 1.637 | 9.03e-05 |  |  |
| LIMLP_13090 (LIC12616/LA1051) |  |  | Hypothetical | 1.578 | 5.23e-10 |  |  |
| LIMLP_13240 (LIC12645/LA1010)** | LEPBI_I0638 |  | Hypothetical | 2.035 | 1.47e-28 |  |  |
| LIMLP_13350 (LIC10866/LA3274) |  |  | Hypothetical | 1.781 | 5.53e-05 |  |  |
| LIMLP_13600** |  |  | Hypothetical | 3.607 | 8.05e-12 |  |  |
| LIMLP_15485 (LIC10451/LA3796) | LEPBI_I0512 |  | Hypothetical | 2.263 | 5.99e-96 |  |  |
| LIMLP_15755 (LEPIC0418/LA0461) | LEPBI_I0512 |  | Hypothetical | 2.804 | 3.35e-19 |  |  |
| LIMLP_16130 (LIC10326/LA0379) | LEPBI_I2376 |  | Hypothetical exported protein | 1.536 | 8.30e-19 |  |  |
| LIMLP_16150 (LIC10322/LA0375) |  |  | Hypothetical | 1.907 | 9.04e-16 |  |  |
| LIMLP_16170 (LEPIC0338/LA0371)** |  |  | Hypothetical | 1.902 | 1.06e-08 |  |  |
| LIMLP_16315 (LIC10291/LA0337)** |  |  | Hypothetical | 2.578 | 1.17e-18 |  |  |
| LIMLP_16745** |  |  | Hypothetical | 2.752 | 1.21e-23 |  |  |
| LIMLP_17465 (LIC13425/LA4279)** |  |  | Hypothetical | 2.277 | 1.93e-02 |  |  |
| LIMLP_18005 (LIC20036/LB050)** | LEPBI_II0173 |  | Hypothetical | 2.015 | 1.18e-43 |  |  |
| LIMLP_18680 (LIC20163/LB207)** | LEPBI_II0256 |  | Hypothetical putative membrane protein | 1.675 | 9.28e-15 |  | 0.569 |
| LIMLP_18815 (LIC20189/LEPIN0211)** | LEPBI_II0190 |  | Hypothetical | 1.735 | 2.10e-08 |  |  |

**S5 Table. Selected up-regulated genes in the *perRAperRB* double mutant.**

Significantly up-regulated genes upon concomitant inactivation of *perRA* and *perRB*.

^a^ Gene numeration is according to Satou et al. (Satou *et al.*, 2015).

^b^ ORF in italic indicates an absence of synteny

^c^ Log_2_FC of significantly differentially-expressed genes (adj. p-value < 0.05) in the *perRA* mutant (M776) (Zavala-Alvarado *et al.*, 2020)

^d^ Log_2_FC of significantly differentially-expressed genes (adj. p-value < 0.05) in the *perRB* mutant (M1474) (this study)

* Up-regulated ORFs upon exposure to 1 mM H_2_O_2_ (adj. p-value < 0.05) (Zavala-Alvarado *et al*., 2020)

** Down-regulated ORFs upon exposure to 1 mM H_2_O_2_ (adj. p-value < 0.05) (Zavala-Alvarado *et al.*, 2020)
