## Supplementary material for "The oxidative stress response of pathogenic *Leptospira* is controlled by two peroxide stress regulators which putatively cooperate in controlling virulence": S6Fig_Zavala_revised_D-20-0288.pdf

**A**

| **Locus**[a] | **Position**[a] | **Reference sequence**[a] | **SNP or Indel** | **Type** | **Change** | **Annotation**[a] | **SNP present in other *L. interrogans* isolates**[b] |
|---|---|---|---|---|---|---|---|
| **LIMLP_01895** | 438255 | GCT | GTT | Non synonymous SNP | A148V | Hybrid Histidine Kinase | no |
| **LIMLP_11570** | 2745296 | ATG | ATGG | Insertion | Frameshift | 3 oxoacyl ACP synthase | 8 isolates |

**B**

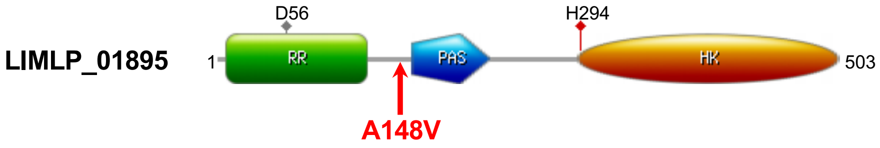


| | | | | | | | |
|---|---|---|---|---|---|---|---|
| WT and *perRB* | …gca<br>…Ala<br>*(145)* | caa<br>Gln | aga<br>Arg | gct<br>Ala | aag<br>Lys | gac<br>Asp | tta…<br>Leu…<br>*(151)* |
| *perRAperRB* | …gca<br>…Ala<br>*(145)* | caa<br>Gln | aga<br>Arg | **gtt**<br>**Val** | aag<br>Lys | gac<br>Asp | tta…<br>Leu…<br>*(151)* |

**S6 Fig**
