## Supplementary material for "The oxidative stress response of pathogenic *Leptospira* is controlled by two peroxide stress regulators which putatively cooperate in controlling virulence": S7Table_ZavalaAlvarado_revised_D-20-0288.docx

| **NC RNA^a^** | **Chromosome/**  **Plasmid** | **Log_2_Fc** | **Adjusted**  **p-value** | **Start-End** | **Overlapping**  **ORF** | **Upstream ORF** | **Downstream**  **ORF** |
| --- | --- | --- | --- | --- | --- | --- | --- |
| **LepncRNA35^#^ (rh753)** | NZ_CP011931.1 | -1.011 | 1.77e-09 | 602707-602840 | NA | LIMLP_02460 | LIMLP_02465 |
| **LepncRNA36** | NZ_CP011931.1 | -1.403 | 5.76e-18 | 611935-611994 | NA | LEPIMA_cI0537 | LIMLP_02490* |
| **LepncRNA87** | NZ_CP011931.1 | -1.187 | 1.15e-12 | 2078455-2078626 | NA | LIMLP_08575 | LIMLP_08580* |
| **LepncRNA89^#§^ (rh2487)** | NZ_CP011931.1 | -1.162 | 1.24e-10 | 2083793-2083898 | LIMLP_08585 | LEPIMA_cI1903 | LIMLP_08590* |
| **LepncRNA109^#§^ (rh3186)** | NZ_CP011931.1 | -1.123 | 2.06e-13 | 2658522-2658603 | NA | LIMLP_11175 | LIMLP_11180** |
| **LepncRNA139** | NZ_CP011931.1 | -1.008 | 1.17e-07 | 3459792-3459865 | NA | LIMLP_14585* | LIMLP_14590 |

**S7 Table. Selected differentially-expressed non-coding RNAs in the *perRB* mutant**

Significantly differentially-expressed predicted ncRNAs in the *perRB* mutant (M1474) with a Log_2_FC cutoff of ±1 and an adjusted p-value cutoff of 0.05.

^a^ Gene numbering is according to Satou *et al.* (2015).

* ORFs significantly down-regulated by RNA-Seq analysis the *perRB* mutant (Log_2_FC cutoff of -0.5 and adj.p-value cutoff of 0.05) (this study).

** ORFs significantly up-regulated by RNA-Seq analysis the *perRB* mutant (Log_2_FC cutoff of 0.5 and adj. p-value cutoff of 0.05) (this study).

^#^ ncRNAs significantly differentially-expressed in the *perRA* mutant (M776) (Zavala-Alvarado *et al*., 2020; the corresponding name is indicated into parenthesis).

^§^ ncRNAs significantly differentially-expressed in the *perRAperRB* mutant (this study).
