## Supplementary material for "The oxidative stress response of pathogenic *Leptospira* is controlled by two peroxide stress regulators which putatively cooperate in controlling virulence": S8Table_ZavalaAlvarado_revised_D-20-0288.docx

| **NC RNA^a^** | **Chromosome/**  **Plasmid** | **Log_2_FC** | **Adjusted**  **p-value** | **Start-End** | **Overlapping**  **ORF** | **Upstream**  **ORF** | **Downstream**  **ORF** | **Log_2_FC**  ***perRA* vs WT** | **Log_2_FC**  ***perRB* vs WT** |
| --- | --- | --- | --- | --- | --- | --- | --- | --- | --- |
| ***Up-regulated*** |  |  |  |  |  |  |  |  |  |
| **LepncRNA3** | NZ_CP011931.1 | 1.798 | 2.36e-13 | 20129-20193 | LIMLP_00080 | LIMLP_00075** | LIMLP_00085** |  |  |
| **LepncRNA30** | NZ_CP011931.1 | 1.339 | 1.03e-05 | 523243-523372 | NA | LIMLP_02195 | LIMLP_02200 |  |  |
| **LepncRNA37** | NZ_CP011931.1 | 1.348 | 1.54e-06 | 635612-635663 | NA | LIMLP_02595** | LIMLP_02600 |  |  |
| **LepncRNA38^#^ (rh859)** | NZ_CP011931.1 | 5.958 | 0.00e-01 | 683753-683935 | NA | LIMLP_02795** | LEPIMA_CI0612 | 2.502 |  |
| **LepncRNA83** | NZ_CP011931.1 | 1.153 | 1.71e-06 | 1925930-1925999 | LIMLP_07855 | LIMLP_07850** | LEPIMA_CI1754 |  |  |
| **LepncRNA109^#§^ (rh3186)** | NZ_CP011931.1 | 1.533 | 6.08e-10 | 2658522-2658603 | NA | LIMLP_11175 | LIMLP_11180** | -0.969 | -1.123 |
| **LepncRNA118** | NZ_CP011931.1 | 1.421 | 3.67e-07 | 2781653-2781712 | NA | LIMLP_11675** | LEPIMA_CI2565 |  |  |
| **LepncRNA127** | NZ_CP011931.1 | 2.406 | 5.13e-52 | 3183156-3183206 | NA | LIMLP_13365 | LIMLP_13370** |  |  |
| **LepncRNA135** | NZ_CP011931.1 | 1.553 | 6.80e-16 | 3366618-3366699 | LIMLP_14160 | LIMLP_14155 | LIMLP_14165 |  |  |
| **LepncRNA140** | NZ_CP011931.1 | 1.172 | 8.04e-04 | 3464676-3464729 | LIMLP_14600 | LIMLP_14595 | LIMLP_14605 |  |  |
| **LepncRNA157** | NZ_CP011931.1 | 1.324 | 2.74e-09 | 3825186-3825268 | NA | LIMLP_16020* | LIMLP_16025** |  |  |
| **LepncRNA166** | NZ_CP011931.1 | 2.690 | 4.87e-143 | 3984145-3984337 | LIMLP_16745** | LIMLP_16740** | LIMLP_16750** |  |  |
| **LepncRNA174** | NZ_CP011932.1 | 1.000 | 3.29e-06 | 24334-24504 | LIMLP_17920* | LEPIMA_CII0022 | LEPIMA_CII0024 |  |  |
| **LepncRNA177** | NZ_CP011932.1 | 1.284 | 3.95e-07 | 125586-125656 | LIMLP_18370** | LIMLP_18365** | LIMLP_18375 |  |  |
| **LepncRNA198^§^** | NZ_CP011933.1 | 1.215 | 5.52e-06 | 34109-34173 | NA | LIMLP_19445** | LEPIMA_p0042 |  | -0.697 |
| **LepncRNA201^§^** | NZ_CP011933.1 | 1.494 | 8.74e-10 | 42542-42606 | NA | LIMLP_19505 | LIMLP_19510 |  | -0.702 |
| **LepncRNA204^§^** | NZ_CP011933.1 | 1.202 | 8.95e-06 | 55785-55880 | NA | LIMLP_19580 | LIMLP_19590 |  |  |
| ***Down-regulated*** |  |  |  |  |  |  |  |  |  |
| **LepncRNA7** | NZ_CP011931.1 | -1.141 | 7.44e-13 | 129161-129223 | LIMLP_00585 | LIMLP_00580 | LIMLP_00590* |  |  |
| **LepncRNA8** | NZ_CP011931.1 | -1.141 | 7.44e-13 | 129339-129401 | LIMLP_00585 | LIMLP_00580 | LIMLP_00590* |  |  |
| **LepncRNA12** | NZ_CP011931.1 | -1.237 | 1.62e-02 | 264596-264849 | NA | LIMLP_01210 | LIMLP_01215* |  |  |
| **LepncRNA34** | NZ_CP011931.1 | -2.067 | 2.76e-95 | 581421-581543 | NA | LIMLP_02390 | LIMLP_02395 |  |  |
| **LepncRNA39** | NZ_CP011931.1 | -3.244 | 0.00e-01 | 695920-696096 | NA | LEPIMA_CI0621 | LIMLP_02840* |  |  |
| **LepncRNA49** | NZ_CP011931.1 | -1.516 | 1.63e-20 | 1030577-1030631 | LIMLP_04255* | LIMLP_04250* | LIMLP_04260 |  |  |
| **LepncRNA65** | NZ_CP011931.1 | -2.510 | 0.00e-01 | 1531021-1531304 | LIMLP_06235 | LIMLP_06230 | LIMLP_06240 |  |  |
| **LepncRNA73** | NZ_CP011931.1 | -1.202 | 9.69e-29 | 1759480-1759649 | NA | LIMLP_07125 | LIMLP_07130* |  |  |
| **LepncRNA74** | NZ_CP011931.1 | -1.112 | 8.35e-20 | 1773996-1774102 | LIMLP_07170* | LIMLP_07165* | LIMLP_07175 |  |  |
| **LepncRNA76** | NZ_CP011931.1 | -1.006 | 5.99e-17 | 1780245-1780377 | LIMLP_07195 | LEPIMA_CI1612 | LIMLP_07200** |  |  |
| **LepncRNA77** | NZ_CP011931.1 | -1.089 | 1.22e-08 | 1809970-1810042 | LIMLP_07330 | LIMLP_07325 | LIMLP_07335 |  |  |
| **LepncRNA79** | NZ_CP011931.1 | -1.633 | 5.85e-41 | 1892073-1892138 | NA | LIMLP_07695** | LIMLP_07700 |  |  |
| **LepncRNA81** | NZ_CP011931.1 | -2.624 | 3.14e-130 | 1902928-1902988 | NA | LIMLP_07735* | LIMLP_07740* |  |  |
| **LepncRNA88^§^** | NZ_CP011931.1 | -1.415 | 1.15e-16 | 2080787-2080896 | NA | LIMLP_08580* | LEPIMA_CI1903 |  | -0.538 |
| **LepncRNA105** | NZ_CP011931.1 | -1.180 | 7.08e-24 | 2612372-2612495 | LEPIMA_CI2416 | LIMLP_10975* | LEPIMA_CI2417 |  |  |
| **LepncRNA119** | NZ_CP011931.1 | -1.394 | 4.41e-26 | 2802847-2803091 | NA | LIMLP_11755* | LIMLP_11760 |  |  |
| **LepncRNA120** | NZ_CP011931.1 | -1.154 | 1.38e-09 | 2900532-2900595 | NA | LIMLP_12145* | LEPIMA_CI2669 |  |  |
| **LepncRNA125** | NZ_CP011931.1 | -2.464 | 5.64e-104 | 3125455-3125516 | NA | LIMLP_13160 | LIMLP_13165 |  |  |
| **LepncRNA128** | NZ_CP011931.1 | -4.413 | 0.00e-01 | 3242298-3242369 | NA | LIMLP_13615 | LIMLP_13620* |  |  |
| **LepncRNA130** | NZ_CP011931.1 | -2.622 | 2.68e-224 | 3252995-3253141 | LIMLP_13675* | LIMLP_13670* | LIMLP_13680 |  |  |
| **LepncRNA133** | NZ_CP011931.1 | -1.122 | 2.06e-02 | 3321048-3321328 | NA | LIMLP_13990* | LIMLP_13995 |  |  |
| **LepncRNA136** | NZ_CP011931.1 | -4.075 | 0.00e-01 | 3379652-3379891 | NA | LIMLP_14225* | LIMLP_14230 |  |  |
| **LepncRNA143** | NZ_CP011931.1 | -2.227 | 6.72e-87 | 3477559-3477624 | NA | LEPIMA_CI3218 | LIMLP_14675 |  |  |
| **LepncRNA144^§^** | NZ_CP011931.1 | -1.015 | 1.75e-17 | 3514242-3514468 | LEPIMA_CI3252 | LIMLP_14810* | LIMLP_14815* |  | -0.507 |
| **LepncRNA145** | NZ_CP011931.1 | -1.087 | 2.72e-11 | 3531452-3531505 | NA | LIMLP_14875 | LEPIMA_CI3268 |  |  |
| **LepncRNA149** | NZ_CP011931.1 | -2.903 | 0.00e-01 | 3697791-3697911 | LIMLP_15435 | LIMLP_15430* | LIMLP_15440 |  |  |
| **LepncRNA155** | NZ_CP011931.1 | -3.354 | 0.00e-01 | 3822733-3822902 | LIMLP_16010 | LIMLP_16005 | LIMLP_16015* |  |  |
| **LepncRNA156** | NZ_CP011931.1 | -1.640 | 5.84e-27 | 3822961-3823020 | NA | LIMLP_16010 | LIMLP_16015* |  |  |
| **LepncRNA162** | NZ_CP011931.1 | -2.066 | 2.38e-74 | 3908300-3908415 | LIMLP_16430 | LIMLP_16425 | LIMLP_16435 |  |  |
| **LepncRNA165^§^** | NZ_CP011931.1 | -1.990 | 8.95e-125 | 3987366-3987576 | LEPIMA_CI3684 | LIMLP_16760 | LIMLP_16765* |  | -0.478 |
| **LepncRNA171** | NZ_CP011931.1 | -1.160 | 9.94e-28 | 4078397-4078517 | NA | LIMLP_17135** | LIMLP_17140 |  |  |
| **LepncRNA175^§^** | NZ_CP011932.1 | -2.120 | 7.13e-55 | 56524-56606 | NA | LIMLP_18085* | LIMLP_18090* |  | -0.720 |
| **LepncRNA179** | NZ_CP011932.1 | -1.910 | 2.15e-100 | 147173-147333 | LIMLP_18455 | LIMLP_18450 | LIMLP_18460** |  |  |
| **LepncRNA180^§^** | NZ_CP011932.1 | -1.091 | 4.23e-14 | 162997-163269 | NA | LIMLP_18530* | LIMLP_18535 |  | -0.530 |
| **LepncRNA182^§^** | NZ_CP011932.1 | -1.158 | 4.41e-23 | 218404-218668 | NA | LIMLP_18735 | LIMLP_18740* |  | -0.397 |
| **LepncRNA194** | NZ_CP011933.1 | -1.373 | 6.09e-10 | 24535-24595 | LIMLP_19390 | LIMLP_19385** | LIMLP_19395* |  |  |
| **LepncRNA208^§^** | NZ_CP011933.1 | -1.058 | 2.49e-16 | 64029-64667 | LIMLP_19625 | LEPIMA_p0089 | LIMLP_19630 |  | -0.374 |
| **LepncRNA210** | NZ_CP011933.1 | -1.029 | 1.43e-05 | 65853-65904 | LIMLP_19635 | LIMLP_19630 | LIMLP_19640 |  |  |

**S8 Table. Selected differentially-expressed non-coding RNAs in the *perRAperRB* mutant**

Significantly differentially-expressed predicted ncRNAs in the *perRAperRB* mutant with a Log_2_FC cutoff of ±1 and an adjusted p-value cutoff of 0.05.

^a^ Gene numbering is according to Satou *et al.* (2015).

* ORFs significantly down-regulated by RNA-Seq analysis in the *perRAperRB* mutant (Log_2_FC cutoff of -0.5 and adj. p-value cutoff of 0.05).

** ORFs significantly up-regulated by RNA-Seq analysis in the *perRAperRB* mutant (Log_2_FC cutoff of 0.5 and adj. p-value cutoff of 0.05).

^#^ ncRNAs significantly differentially-expressed in the *perRA* mutant (M776), as determined by Zavala-Alvarado *et al.*, (2020) with the corresponding name indicated into parenthesis.

^§^ ncRNAs significantly differentially-expressed in the *perRB* mutant (M1474) (this study).
