## Supplementary material for "The oxidative stress response of pathogenic *Leptospira* is controlled by two peroxide stress regulators which putatively cooperate in controlling virulence": S9Table_ZavalaAlvarado_revised_D-20-0288.docx

**Strains	Description^a^	Tn insertion site^b^	Source**

____________________________________________________________________________________________________________________

*Leptospira interrogans*	Wild-type strain (WT)

Serovar Manilae, strain L495

*perRA* (M766)	*Himar1* Tn insertion in LIMLP_10155 (2427923-2428360)	2427985	Murray *et al.* (2009)	*perRA::Km^R^*	Lo *et al.* (2010)

Resistance to kanamycin

*perRB* (Man1474)	*Himar1* Tn insertion in LIMLP_05620 (1386251-1386688)	1386423	This study	*perRB::Km^R^*

Resistance to kanamycin

*perRAperRB*	allelic exchange of LIMLP_10155 in the M1474 mutant	1386423	This study

Δ *perRA, perRB::Km^R^*

Resistance to kanamycin and spectinomycin

*perRB*^+^*^perRB^*	Man1474 mutant trans-complemented with the LIMLP_05620 ORF	This study

contains the pNB139 plasmid

Resistance to kanamycin and spectinomycin

Π1 (Δ*thyA*)	*Escherichia coli* replicative strain

β2163 (Δ*dapA*)	*Escherichia coli* conjugative strain	Demarre *et al.* (2005)

^a^ Gene name is according to *Leptospira interrogans* serovar Manilae strain UP-MMC-NIID-LP genome (Satou *et al.* (2015)).

^b^ Tn position is according to *Leptospira interrogans* serovar Manilae strain UP-MMC-NIID-LP genome (Satou *et al.* (2015)).

**S9 Table. Strains used in this study**
