## Supplementary material for "The oxidative stress response of pathogenic *Leptospira* is controlled by two peroxide stress regulators which putatively cooperate in controlling virulence": S10Table_ZavalaAlvarado_revised_D-20-028.docx

**Plasmids	Description** ^a^	**Source**

pMaORI	replicative and conjugative vector for *Leptospira*	Pappas et al. (2015)

pNB139	pMaORI containing the LIMLP_05620 (*perRB*) ORF	This study

Promoter: 200 bp upstream region

Resistance to spectinomycin

pKΔperRA	suicide plasmid containing a spectinomycin resistance cassette	This study

flanked by the LIMLP_10155 (*perRA*) ORF

^a^ Gene name is according to *Leptospira interrogans* serovar Manilae strain UP-MMC-NIID-LP genome (Satou *et al.* (2015)).

**S10 Table. Plasmids used in this study**
