## Supplementary figures and images for "The oxidative stress response of pathogenic *Leptospira* is controlled by two peroxide stress regulators which putatively cooperate in controlling virulence"

### S1Fig_ZavalaAlvarad_revised_D-20-0288.pdf

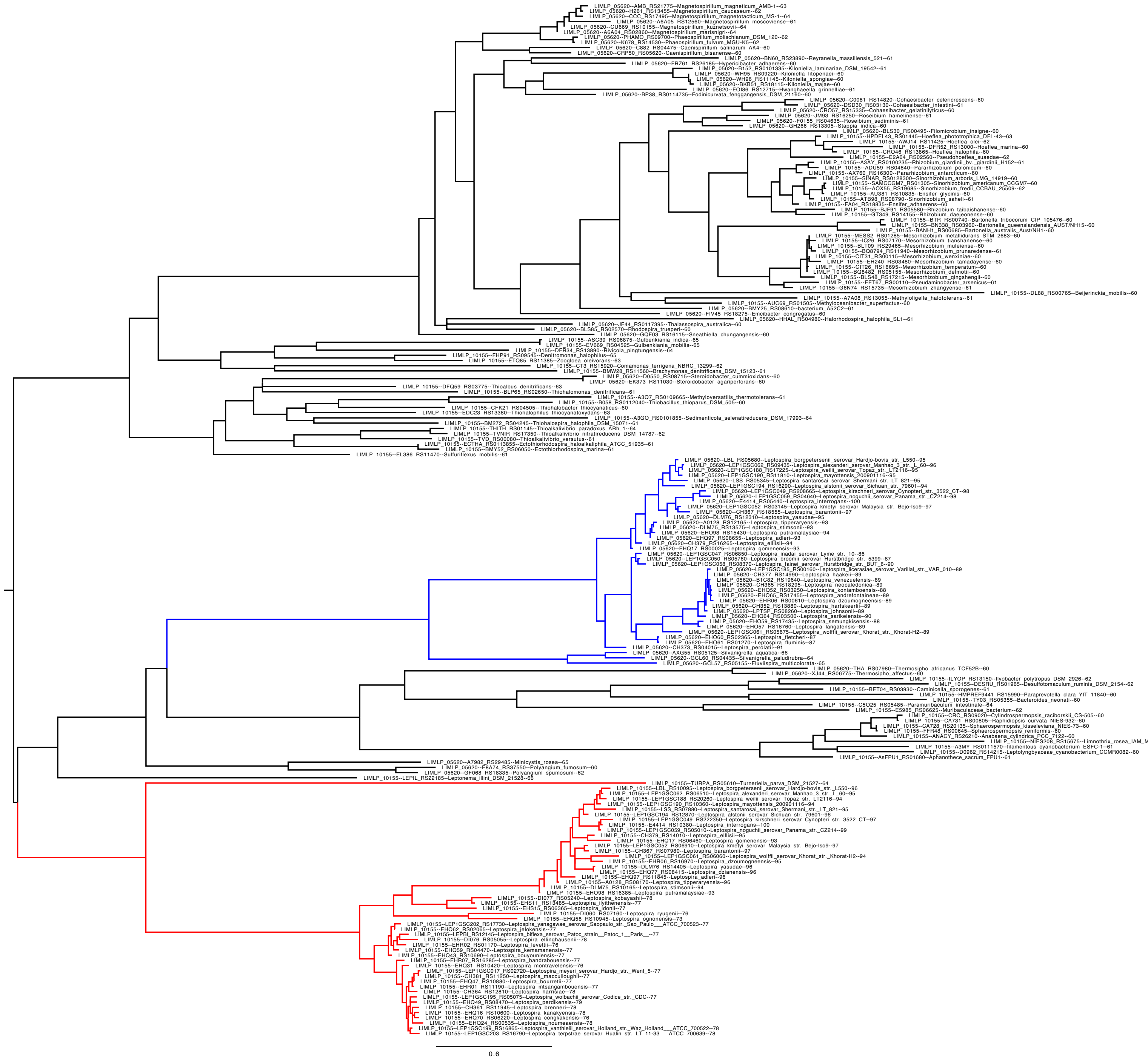


**S1 Fig**

### S2Fig_ZavalaAlvarado_revised_D-20-0288.pdf

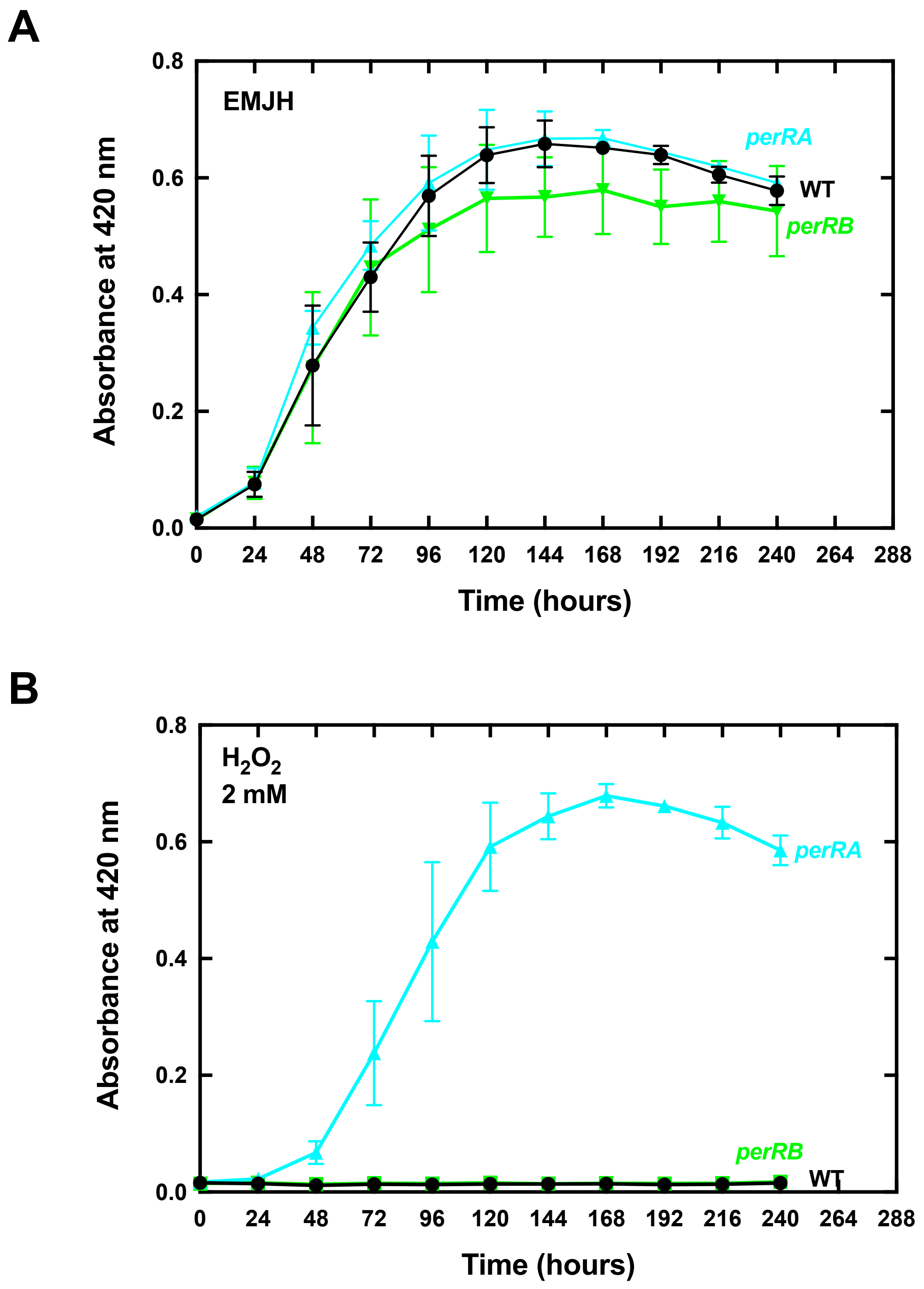

### S3Fig_ZavalaAlvarado_revised_D-520-0288.pdf

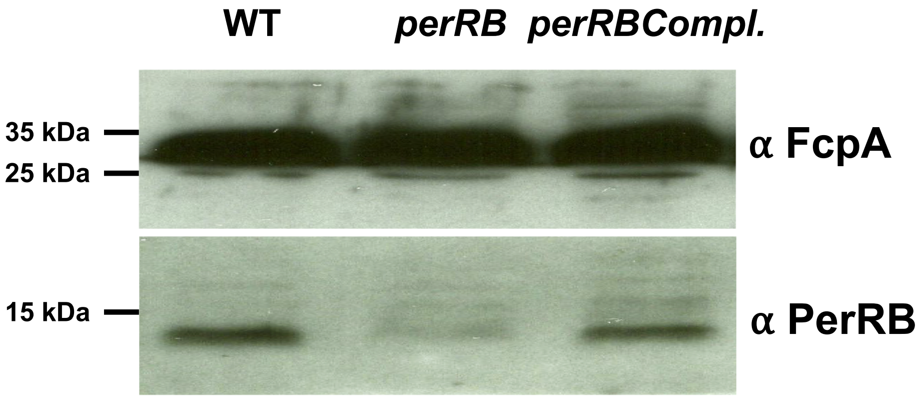

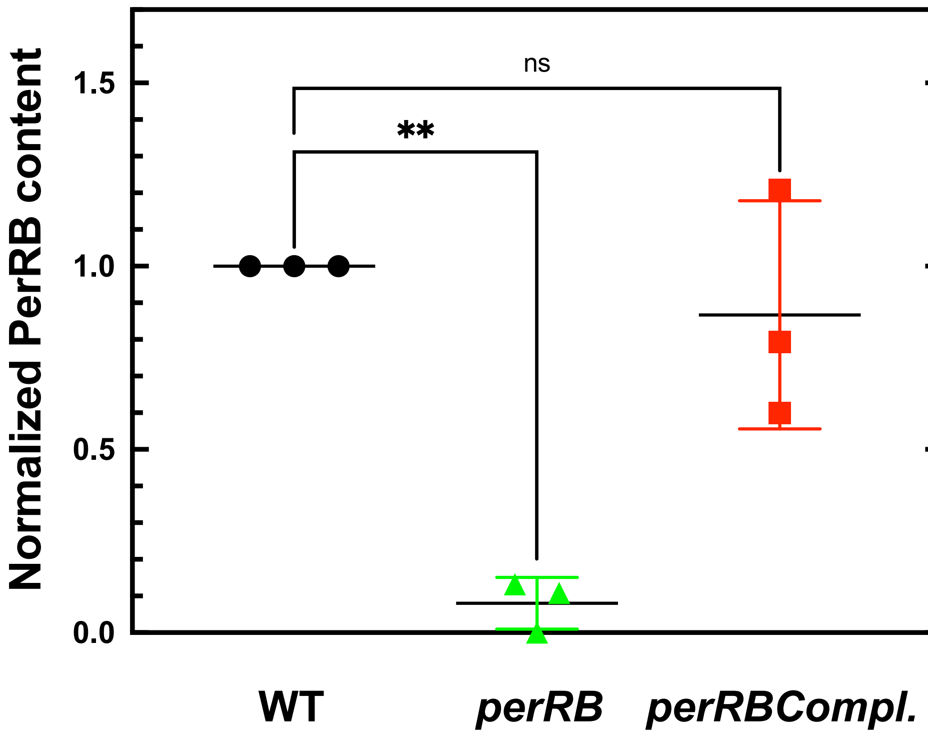

### S4Fig_Zavala2_revised_D-20-0288.pdf

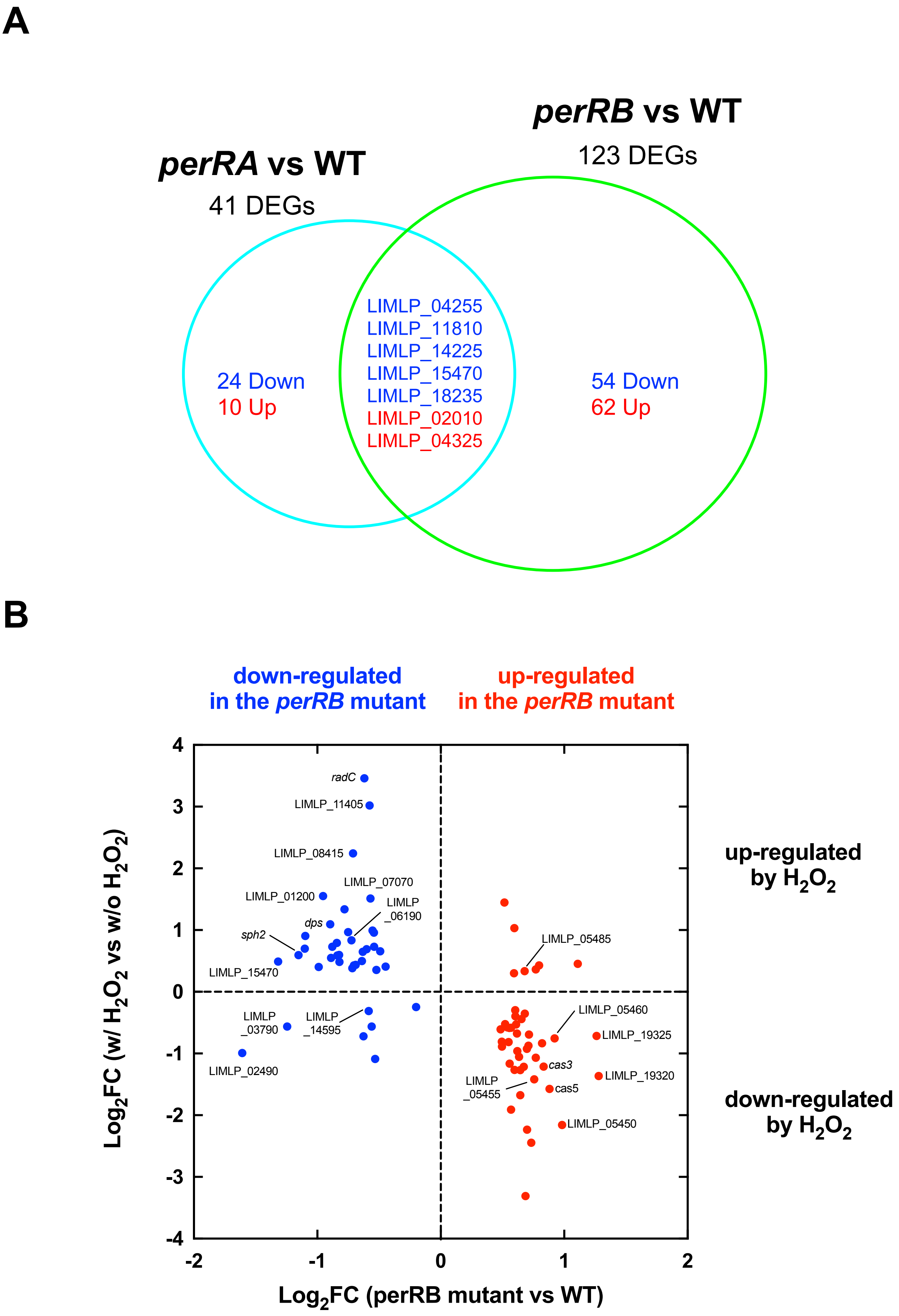

### S5Fig_Zavala2_revised_D-20-0288.pdf

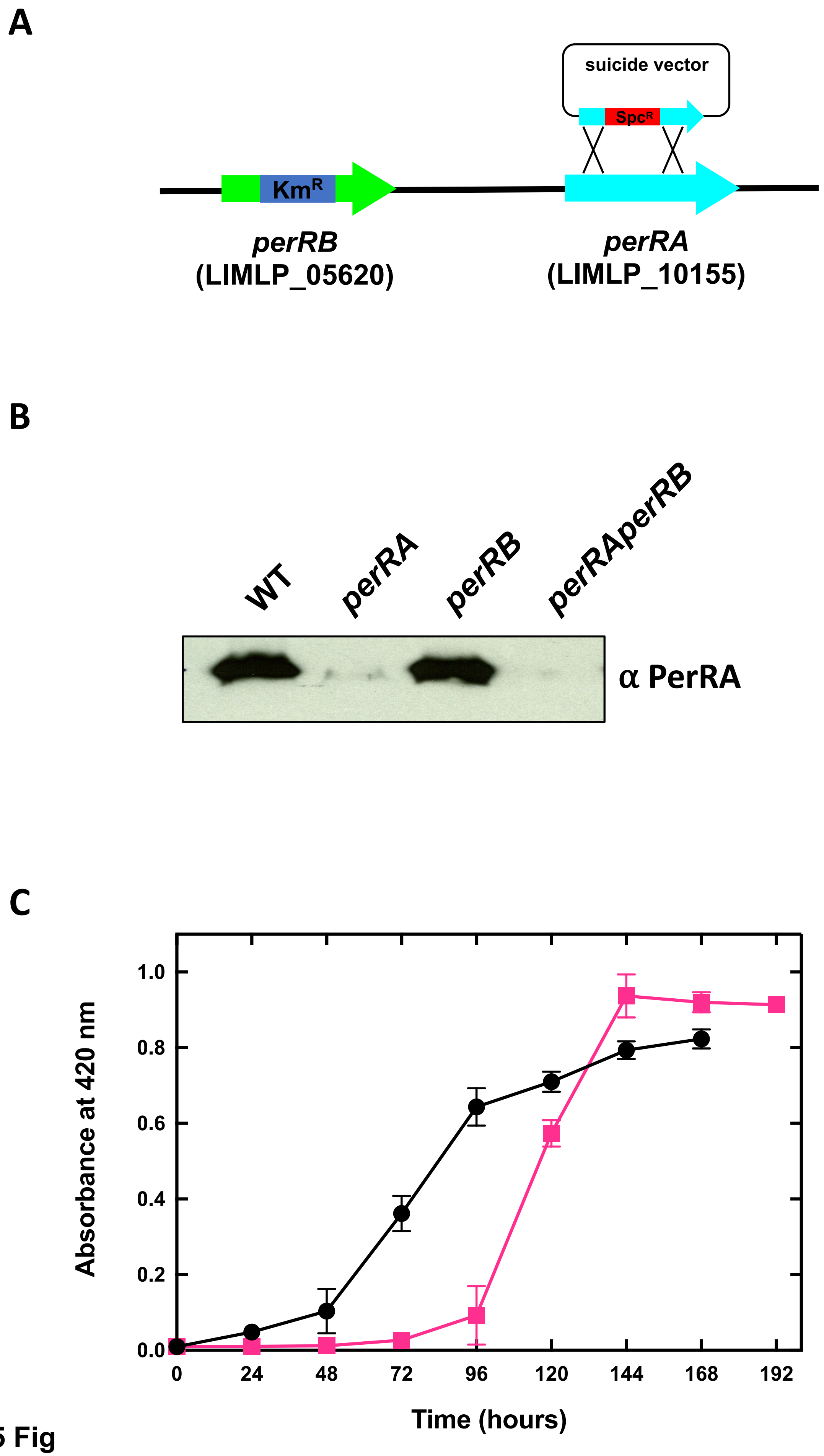

### S7Fig_Zavala2_revised_D-20-0288.pdf

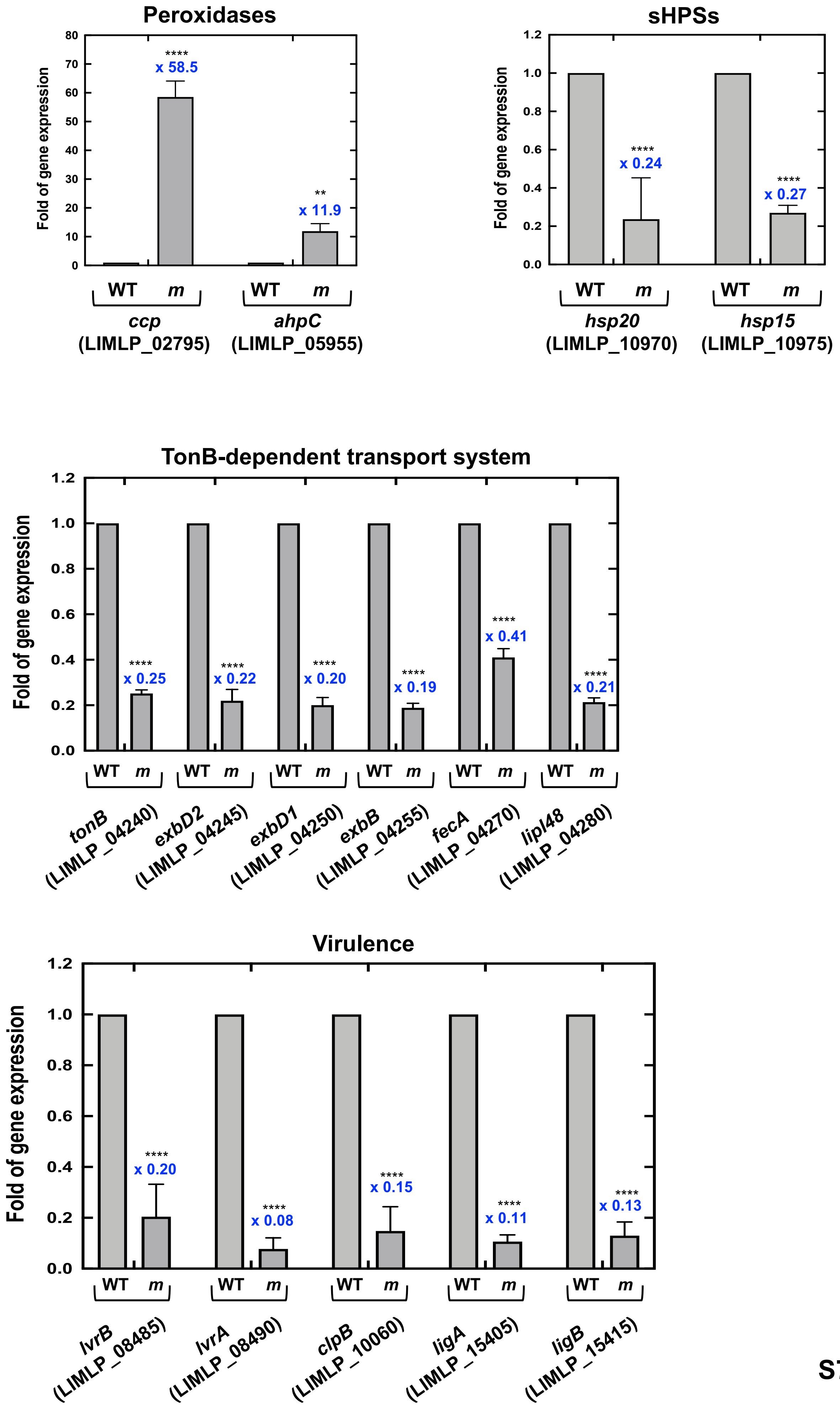

### S8Fig_ZavalaAlvarado_revised_D-20-0288.pdf

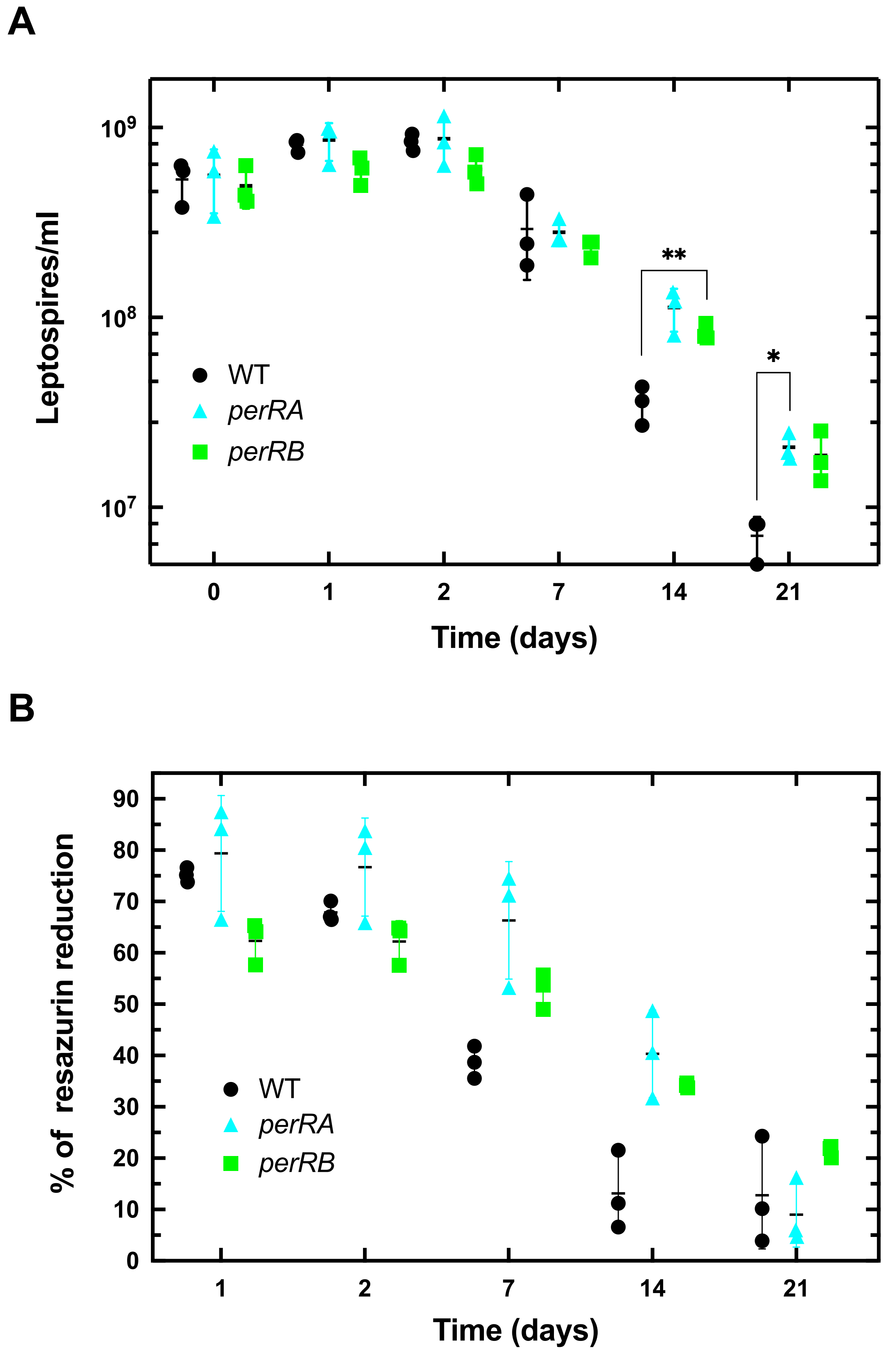
